# Wnt/β-catenin signaling promotes zebrafish osteoblast dedifferentiation by *wnt10a*-mediated inhibition of NF-κB

**DOI:** 10.64898/2025.12.29.696582

**Authors:** Hossein Falah Mohammadi, Denise Posadas Pena, Dila Gülensoy, Ivonne Sehring, Gilbert Weidinger

**Affiliations:** Institute of Biochemistry and Molecular Biology, Ulm University, Albert-Einstein-Allee 11, 89081 Ulm, Germany

## Abstract

Dedifferentiation of mature cells is an essential mechanism of source cell formation for regeneration of many systems, including the zebrafish fin. Here, we use bulk and single cell RNASeq to show that osteoblast injury responses involve rapid and extensive transcriptional reprogramming, yielding a cell state that shares characteristics with embryonic osteoblasts, but also expresses many regeneration-specific genes. One gene characterizing this state is the canonical Wnt ligand *wnt10a*. Using genetic and transgenic perturbations, we demonstrate that *wnt10a*-dependent Wnt/β-catenin signaling cell-autonomously induces osteoblast dedifferentiation. Loss of *wnt10a* or of Wnt/β-catenin activity blocks the dedifferentiation program, whereas *wnt10a* overexpression enhances dedifferentiation and is sufficient to induce it even without injury. Wnt/β-catenin signaling promotes dedifferentiation by suppressing NF-κB activity, placing it upstream of known cues whose loss causes dedifferentiation. Notably, *wnt10a* overexpression also stimulates cardiomyocyte dedifferentiation during zebrafish heart regeneration, revealing a conserved role in activating source-cell formation during regeneration.

## Introduction

Dedifferentiation of mature adult cells into proliferative progenitors is an important mode of how source cells for regeneration are generated. We define dedifferentiation as the (partial) reversal of the specialization of a differentiated cellular phenotype, which is usually accompanied by cell cycle entry and the acquisition of characteristics of progenitor cells. For some cell types, e.g. cardiomyocytes in the regenerating adult heart of fish or salamanders, or of neonatal mice, it is the only available means for regeneration, since (adult) stem cells for cardiomyocytes do not exist ^1,2^. Other cell types, like newt muscle fibers during limb regeneration, employ dedifferentiation despite the existence of stem cells ^3^, while in several mammalian systems, dedifferentiation of mature or transit amplifying cells serves as a backup means for source cell production in case the normal mode of stem-cell driven regeneration fails ^4,5^. Thus, naturally occurring dedifferentiation represents a fascinating biological phenomenon that in many systems is essential for robust regeneration. A thorough understanding of the underlying molecular mechanisms is not only of great biological interest, but might also aid in the development of regenerative therapies.

Zebrafish, like most teleost fish, can efficiently and robustly regenerate their fins after partial amputation, including bone forming the fin rays, which stands in stark contrast to the inability of adult mammalian limbs to regenerate. Mature osteoblasts dedifferentiate in response to fin injuries, and migrate towards the injury to provide source cells for regeneration ^6–10^. In contrast, current evidence suggests that differentiated osteoblasts do not participate in bone fracture repair in mammals, which seems to be entirely driven by skeletal stem cells ^11^. While as-yet poorly characterized alternative cellular sources for bone regeneration also exist in fins ^12,13^, it is plausible that the ability of fish to recruit mature differentiated osteoblasts as source cells for bone regeneration is at least partly responsible for the high regenerative abilities of fins.

The molecular mechanisms regulating osteoblast dedifferentiation are only beginning to emerge. NF-ĸB signaling is active in mature osteoblasts and needs to be downregulated for osteoblast dedifferentiation to occur ^14^. NF-ĸB signaling appears to act upstream of retinoic acid signaling, which likewise interferes with dedifferentiation ^14,15^. The only other regulatory mechanism that has been shown to at least indirectly impinge on osteoblast dedifferentiation is a metabolic switch from mitochondrial oxidation to glycolysis, which is necessary for formation of pre-osteoblasts ^16^. Thus, significant gaps remain in our understanding of the molecular mechanisms driving osteoblast dedifferentiation.

Wnt/β-catenin signaling plays essential roles in the regeneration of many systems, from planaria to mammals, and is also required for zebrafish fin regeneration ^17–21^. When Wnt/β-catenin signaling is blocked, fin stumps heal the epidermal wound, but subsequent formation of a regenerate does not occur ^21,22^. While this hints at an early requirement of Wnt signaling in the stump for formation of source cells for regeneration, such a function has not been characterized. Yet, at later stages of regeneration, Wnt/β-catenin signaling again plays important roles for outgrowth of the regenerate. In particular, Wnt signaling organizes proliferation of progenitor cells in the regenerate and differentiation of osteoblasts from osteogenic progenitors, acting upstream of many other signaling pathways ^19–21^;.

Here we find that injury-induced osteoblast dedifferentiation is characterized by attainment of a specific regenerative transcriptional state that resembles embryonic osteoblasts, but also includes expression of many regeneration-specific transcripts, including *wnt10a*. Using transgenics and mutants, we show that Wnt/β-catenin signaling, activated by *wnt10a*, acts within osteoblasts to promote their dedifferentiation. Intriguingly, *wnt10a* overexpression is sufficient to induce osteoblast dedifferentiation even in the absence of fin injury, and *wnt10a* can also promote the dedifferentiation of cardiomyocytes during zebrafish heart regeneration. Epistasis experiments show that Wnt/β-catenin signaling acts upstream of NF-ĸB signaling in the regulation of osteoblast dedifferentiation. Our findings reveal that Wnt/β-catenin signaling does not only act as master regulator of regenerative growth and patterning, but is also key for the production of source cells for regeneration.

## Results

### Transcriptomics reveals extensive transcriptional changes associated with osteoblast dedifferentiation

Osteoblasts dedifferentiate in response to injuries of fin and skull bones in zebrafish, and migrate towards the injury to provide source cells for regeneration ^6,8,10,14,16^. Dedifferentiation occurs within 1 day post fin amputation (dpa) in a graded fashion across a zone encompassing several hundred micrometers proximal of the amputation plane (**Figure 1A**) ^6,10^. In the segmented fin ray, dedifferentiation does not only occur in the amputated segment (segment 0), but also in the adjacent non-injured segment -1 (**Figure 1A).** While we and others have previously shown that dedifferentiating osteoblasts downregulate several markers of differentiation, including *bglap* and *entpd5a*, and upregulate progenitor markers, a more comprehensive picture of transcriptional changes associated with osteoblast dedifferentiation has been missing ^6,8,10,14,16^. Thus, we performed RNASeq of osteoblasts isolated from the *Ola.Bglap.1:EGFP^hu^*^4008^ transgenic line (abbreviated *bglap:GFP* ^10^). GFP labels a subset of the differentiated osteoblasts, located in the centers of each bony segment (**Figure 1B**). While transcription of the endogenous zebrafish *bglap* gene and of the transgenic *gfp* RNA is lost within hours after fin amputation in dedifferentiating osteoblasts, the GFP protein persists for several days in these cells ^6,10,14^. Thus, the *bglap:GFP* transgenic line can be used to isolate differentiated osteoblasts during homeostasis and dedifferentiated osteoblasts after fin amputation (**Figure 1B**). RT-qPCR showed that GFP+ cells FAC-sorted from homeostatic, non-injured fins were strongly enriched for markers of differentiated osteoblasts compared to the GFP– fraction, and deriched for genes expressed in the epidermis and in fibroblasts (**Figure 1C, Supplementary Fig. 1A**). Bulk RNA-Seq of GFP+ and GFP– cells isolated from non-injured fins confirmed that GFP+ cells represent mature osteoblasts that are enriched in genes annotated to participate in ossification and bone mineralization including *bglap, bglapl*, *entpd5a*, and *col10a1a* (**Figure 1D**, **Supplementary Fig. 1B-E**).

**Figure 1.**
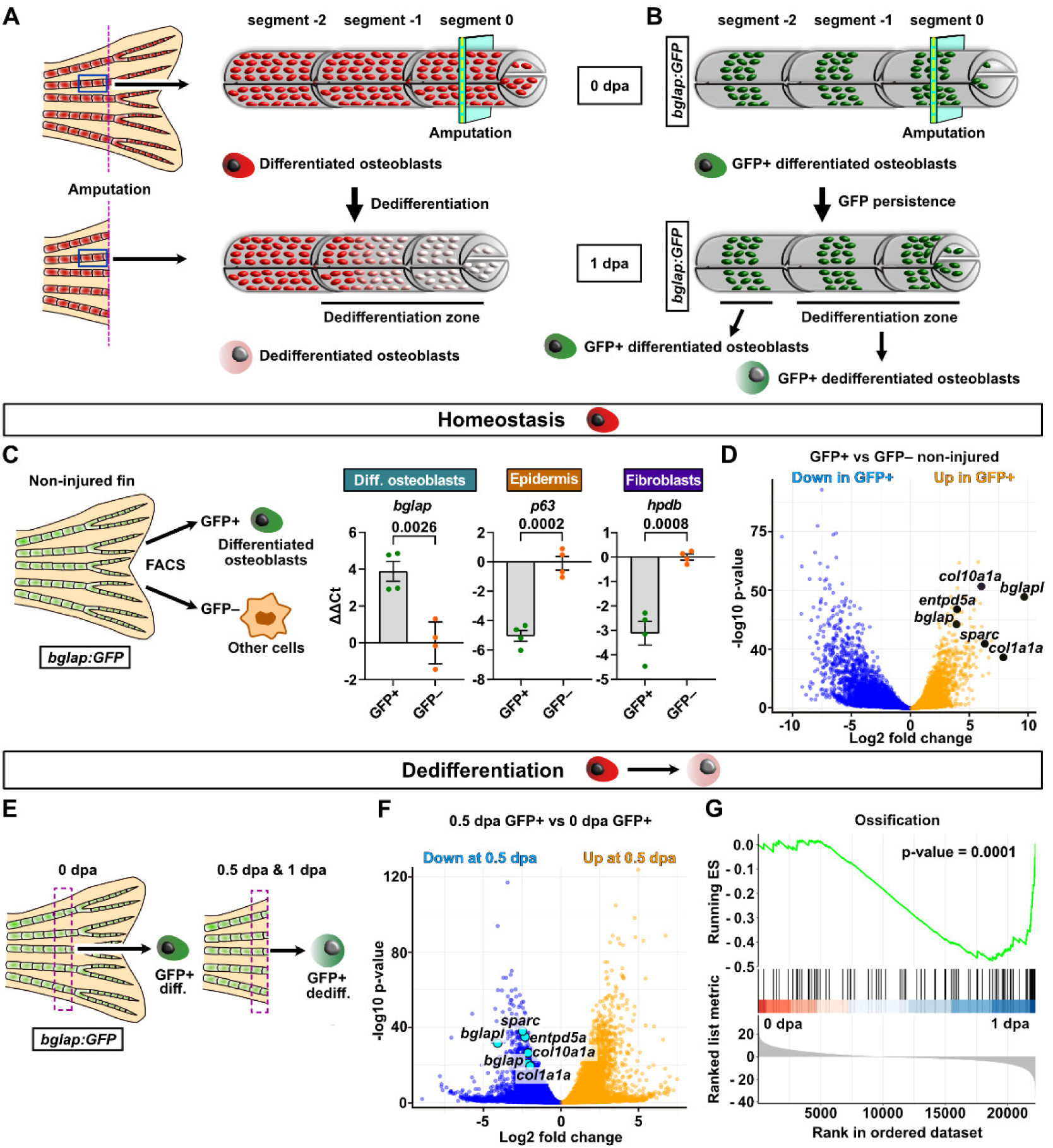
RNASeq of sorted fin osteoblasts reveals extensive transcriptional changes during dedifferentiation. **A)** In response to amputation of the zebrafish caudal fin, osteoblasts dedifferentiate within 1 day post amputation (dpa) in a zone encompassing the amputated bony segment (segment 0) and the adjacent segment -1. **B)** GFP labels a subset of differentiated osteoblasts in *bglap:GFP* transgenic fish. After shutdown of transgene transcription in dedifferentiating osteoblasts, GFP protein persists, allowing for sorting of dedifferentiating osteoblasts at 1 dpa. **C)** RT-qPCR on GFP+ and GFP– cells sorted from non-injured homeostatic *bglap:GFP* fins reveals enrichment of the differentiated osteoblasts marker *bglap* in the GFP+ fraction, and little expression of markers of the epidermis (*p63*) or fibroblast (*hpdb*). ΔΔCt values are shown relative to the mean of the GFP– samples. nE (biological replicates) = 4, nA (animals) = 10 per replicate. Error bars, mean ± SEM. Numbers indicate p-value. Two tailed Student’s t-test. **D)** Volcano plot of bulk RNASeq data showing genes differentially expressed between GFP+ and GFP–fractions from non-injured fins. **E)** GFP+ cells were sorted from the dedifferentiation zone of *bglap:GFP* fins at 0.5 and 1 dpa and from the same region at 0 dpa. **F)** Volcano plot of differentially expressed genes between GFP+ osteoblasts at 0.5 dpa and 0 dpa reveals that known differentiation genes are downregulated at 0.5 dpa. **G)** Gene set running enrichment score plot indicating the downregulation of ossification -related genes based on the GO terms “biological process” at 1 dpa compared to 0 dpa.

Having validated our experimental approach, we FAC-sorted differentiated osteoblasts isolated immediately after fin amputation (0 day post amputation, dpa) and dedifferentiating osteoblasts at 12 hours post amputation (0.5 dpa) and at 1 dpa from the same region of *bglap:GFP* transgenic fins (the “dedifferentiation zone”, encompassing segments 0 and -1 in amputated fins, **Figure 1A, E**) and subjected them to bulk RNA-Seq (**Supplementary Fig. F, G**). We found that extensive transcriptional changes occur in GFP+ cells during dedifferentiation with almost 29% of the transcriptome (> 7000 genes) up- or downregulated relative to 0 dpa at 0.5 dpa (**Figure 1F, Supplementary Fig. 1H, J**) and 44% of all transcripts (> 11000 genes) at 1 dpa (**Supplementary Fig. 1I, J**). By 0.5 dpa, known osteoblast differentiation markers were already downregulated (**Figure 1F**), and most were even more strongly downregulated by 1 dpa (**Supplementary Fig. 1K)**. Gene sets related to ossification and osteoblast differentiation were negatively enriched by 1 dpa (**Figure 1G, Supplementary Fig. 1L**). We conclude that osteoblasts react to fin amputation rapidly with extensive transcriptional reprogramming.

### Identification of dedifferentiation gene signatures

To validate the RNA-Seq data and to identify previously unknown genes associated with zebrafish osteoblast differentiation, we selected genes that were more strongly expressed in GFP+ vs GFP– cells isolated from non-injured fins and were also downregulated in dedifferentiating GFP+ osteoblasts at 0.5 dpa and 1 dpa (**Figure 2A**). RT-qPCR confirmed osteoblast-specific expression of known differentiation markers during homeostasis and of a number of genes not previously recognized as being enriched in differentiated zebrafish osteoblasts, including *bmp8a*, *cadm1b*, *ccn4b*, *cd81b, hhipl2*, and *sema3aa*, (**Figure 2B**). Importantly, all of these genes were also downregulated in sorted dedifferentiating osteoblasts at 0.5 dpa (**Figure 2C**) and at 1 dpa (**Supplementary Fig. 2A),** except for *col1a1a*, *col10a1a* and *hhipl2*, which were only transiently down at 0.5 dpa. In situ hybridization confirmed osteoblast-specific expression and downregulation of *sparc*, *ccn4b*, *bgnb*, and *clec11a* at 0.5 dpa in segment 0 and for some genes also in segment -1 (**Figure 2D** and Error! Reference source not found.**2B**).

**Figure 2.**
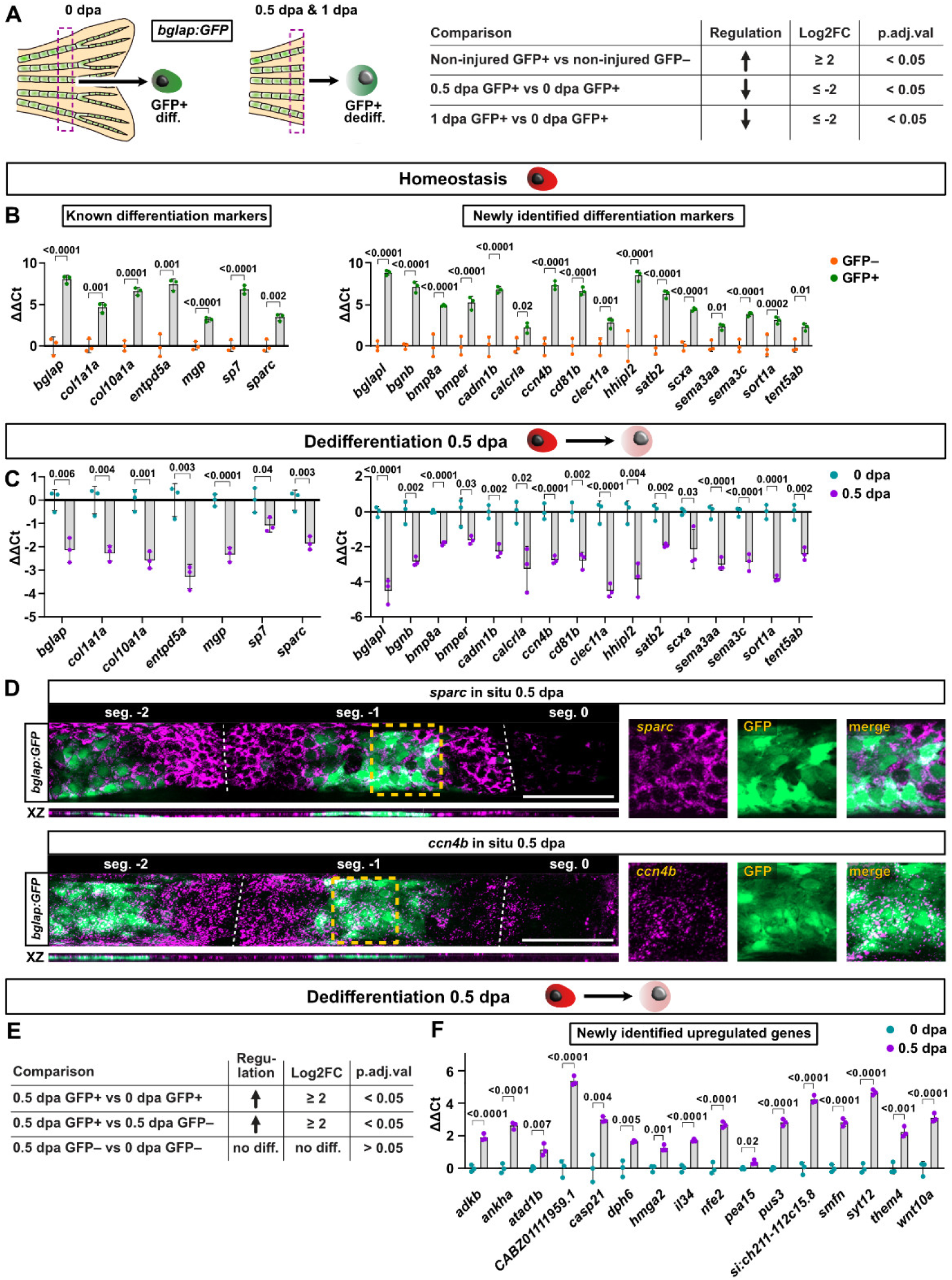
Identification of novel markers of the osteoblast differentiated state that are downregulated during dedifferentiation. **A)** Criteria used to filter the RNASeq data for genes specific to differentiated osteoblasts that are downregulated during dedifferentiation. **B)** RT-qPCR confirms enrichment of known genes specific to differentiated osteoblasts (left), and of newly identified genes (right) in the GFP+ fraction of non-injured *bglap:GFP* fins compared to the GFP– fraction. nE (biological replicates) = 3, nA = 15 per replicate. **C)** RT-qPCR confirms downregulation of the same genes shown in (B) in *bglap:GFP*+ samples at 0.5 dpa compared to 0 dpa. nE (biological replicates) = 3, nA = 15 per replicate. **D)** HCR in situ hybridization confirms specific expression of *sparc* and *ccn4b* in osteoblasts (identified by location in the XZ view, and by co-localization with *bglap:GFP*) and downregulation in segment 0 at 0.5 dpa. nE (independent experiments) = 1, nA = 6, nR (rays) = 6. Dashed line, joints. Scale bar, 100 µm. **E)** Filtering criteria used to identify genes that are upregulated in dedifferentiating osteoblasts. **F)** RT-qPCR confirms upregulation of selected genes fulfilling the criteria in (E) in *bglap:GFP*+ samples at 0.5 dpa compared to 0 dpa. nE (biological replicates) = 3, nA = 15 per replicate. **(B, C, F)** Data are presented as mean values. ΔΔCt values are shown relative to the mean of the GFP–samples (B) or the 0 dpa samples (C, F). Error bars, mean ± SEM. Two tailed Student’s t-test.

To identify genes that are specifically upregulated in dedifferentiating osteoblasts, we intersected our transcriptomics data using the criteria specified in **Figure 2E**. We picked several genes for validation based on strength of regulation and predicted biological function. RT-qPCR confirmed upregulation of these genes in sorted *bglap:GFP*+ osteoblasts at 0.5 and 1 dpa, with the exception of *nfe2* and *pea15,* which were only transiently upregulated at 0.5 dpa (**Figure 2F, Supplementary Fig. 3A**). Validated genes hint at cell-autonomous functions of inflammatory signaling (*il34, casp21, si:ch211-112c15.8* – a homolog of *Tnfrsf1a*), (oxidative) stress response (*nfe2, atad1b, smfn*), adenosine signaling (*adkb*), and protein biosynthesis quality control (*dph6, pus3*) in dedifferentiating osteoblasts. In situ hybridization confirmed upregulation of *nfe2* and *hmga2* specifically in osteoblasts close to the amputation plane at 0.5 dpa (**Supplementary Fig. 3B**).

To characterize different osteoblast cell states during the process of dedifferentiation in more detail, we turned to single cell RNASeq. We enriched for osteoblasts using FACS with the Zns5 antibody **(Supplementary Fig. 4A)**, which recognizes an uncharacterized cell-surface antigen that is expressed on all osteoblasts in zebrafish fins, including populations that are not fully differentiated, which are for example located at the joints between bony segments, plus some other mesenchymal cell types. RT-qPCR on cells isolated from homeostatic, non-injured fins confirmed that the Zns5+ population was enriched for markers of differentiated, committed, and pre-osteoblasts, but deriched for epidermal cells (**Figure 3A**). Next, we isolated Zns5+ cells at 0 dpa, 6 hours post amputation (hpa), 0.5 dpa and 1 dpa (**Figure 3B**). Analysis of the combined data from all stages revealed 12 cell clusters, little contamination with epidermis and macrophages, and several clusters representing osteoblasts, including differentiated osteoblasts, based on known differentiation markers and those newly identified by our bulk RNA-seq (**Figure 3C, D, Supplementary Fig. 4B, C**). At 0 dpa, the Zns5+ population also included three clusters that we assume represent less differentiated osteoblast states located at the joints between bony segments, plus fibroblasts and two clusters whose identity and location in the fin warrants future studies, which we termed “mesenchyme 1 and 2” (**Figure 3E**). Importantly, after fin amputation, three new clusters rapidly appeared by 6 hpa, one of unknown character, one that seems to be related to the mesenchymal clusters (“mesenchyme 3”) and one that we assume represents dedifferentiated osteoblasts based on enrichment in genes like *nfe2* and *syt12* that we have shown above to be upregulated during dedifferentiation (**Figure 3D, E**). The “unknown” and “dedifferentiated osteoblasts” clusters increased in prominence through 0.5 and 1 dpa (**Figure 3E**). Comparison of the “differentiated osteoblasts” and the “dedifferentiated osteoblasts” clusters confirmed that all genes that we have shown by RT-qPCR to be downregulated during dedifferentiation were enriched in the “differentiated osteoblasts” cluster, while all but one gene that we had found to be upregulated were enriched in the “dedifferentiated osteoblasts” cluster (**Figure 3F, G, Supplementary Figure 5A, B**). Thus, bulk and scRNASeq nicely complement each other and identify a high-confidence set of dedifferentiation-associated genes.

**Figure 3.**
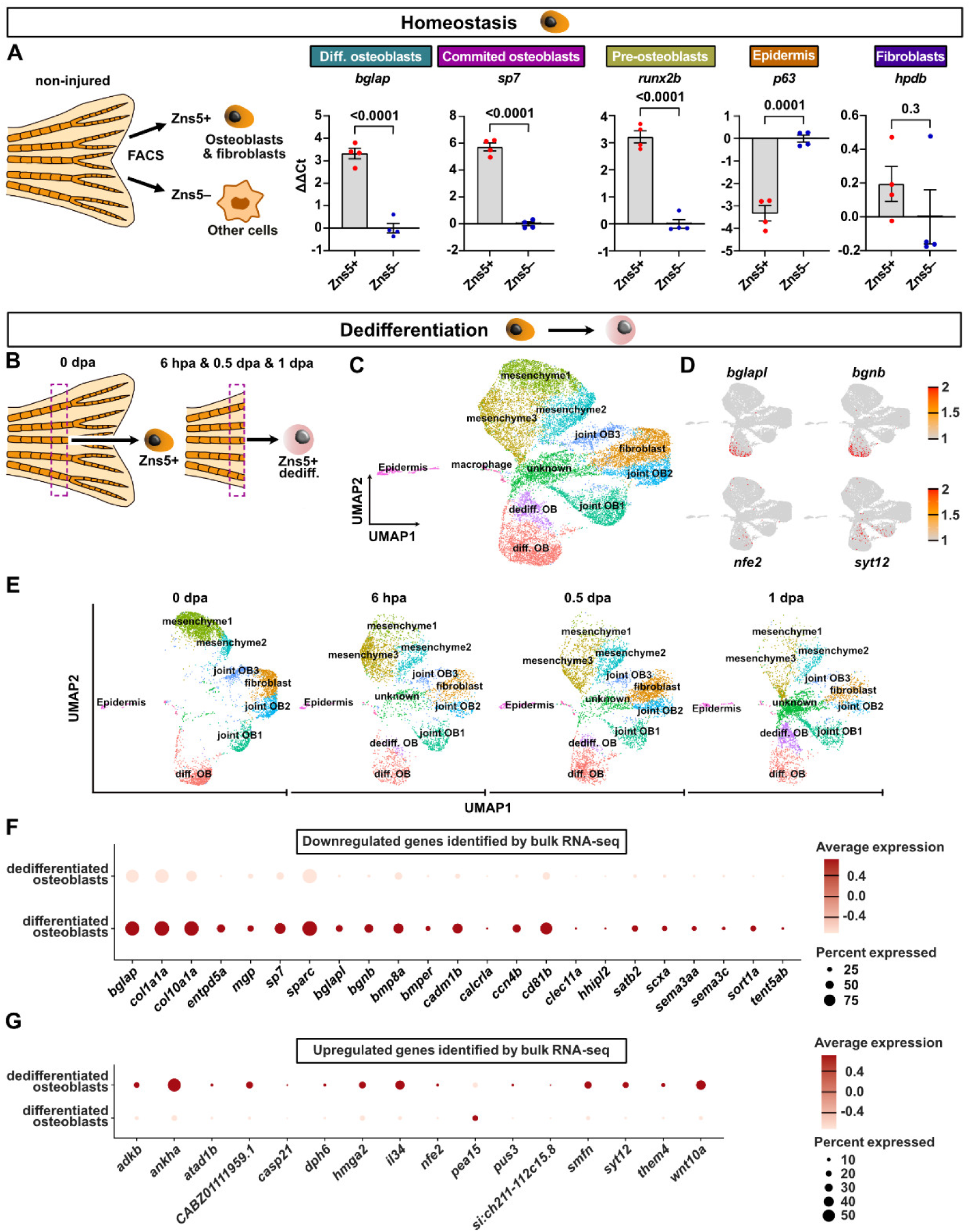
scRNASeq identifies osteoblast state transitions induced by fin amputation. **A)** RT-qPCR analysis of Zns5+ cells sorted from non-injured fins shows that this population includes osteoblasts of various maturation states, plus fibroblasts. nE (biological replicates) = 4, nA = 10 per replicate. ΔΔCt values are shown relative to the mean of the Zns5– samples. Error bars, mean ± SEM. Two tailed Student’s t-test. **B)** Zns5+ cells sorted from the dedifferentiation zone at 6 hpa, 0.5 dpa and 1 dpa and from the same region at 0 dpa were subjected to scRNA sequencing. **C)** UMAP embedding of scRNAseq data of 38296 cells of all above-mentioned time points combined. Cells are colored by cluster. **D)** Feature plots showing the expression of markers of differentiated osteoblasts (top) and of genes upregulated during dedifferentiation (bottom). **E)** UMAP embedding showing cluster distribution separated by time point. **F)** Dot plot comparing downregulated genes identified by bulk RNA sequencing in differentiated and dedifferentiated osteoblast clusters (clusters of cells from all time-points combined). **G)** Dot plot comparing upregulated genes identified by bulk RNA sequencing in differentiated and dedifferentiated osteoblast clusters.

### Osteoblasts dedifferentiate into a regenerative state that partly resembles embryonic osteoblasts

We next asked whether dedifferentiated osteoblasts resemble embryonic osteoblasts, or represent a regeneration-specific cell state. To this end, we compared our datasets to a published bulk transcriptome of *sp7*+ osteoblasts sorted from 4 days post fertilization (dpf) embryos ^23^. Unsupervised clustering of the overall bulk RNASeq expression patterns revealed that dedifferentiating 0.5 and 1 dpa fin osteoblasts were more similar to 0 dpa fin osteoblasts than to 4 dpf embryonic osteoblasts (**Figure 4A**). Yet, a substantial number of genes whose expression changes in osteoblasts with dedifferentiation (0.5 dpa and 1 dpa fin vs 0 dpa fin osteoblasts) is also differentially expressed between embryonic and 0 dpa fin osteoblasts (e.g. 46% of downregulated genes at 0.5 dpa are also more weakly expressed in embryonic osteoblasts, 28% of genes upregulated at 1 dpa are also higher in embryonic osteoblasts, **Supplementary Fig. 6A**). Likewise, heatmap of those genes that we have confirmed by RT-qPCR to be up- and downregulated in dedifferentiating osteoblasts revealed many similarities with the expression profile of embryonic osteoblasts (**Supplementary Fig. 6B**). We conclude that osteoblast injury responses result in dedifferentiation into a state that is not identical to, but shares many similarities with embryonic osteoblasts.

**Figure 4.**
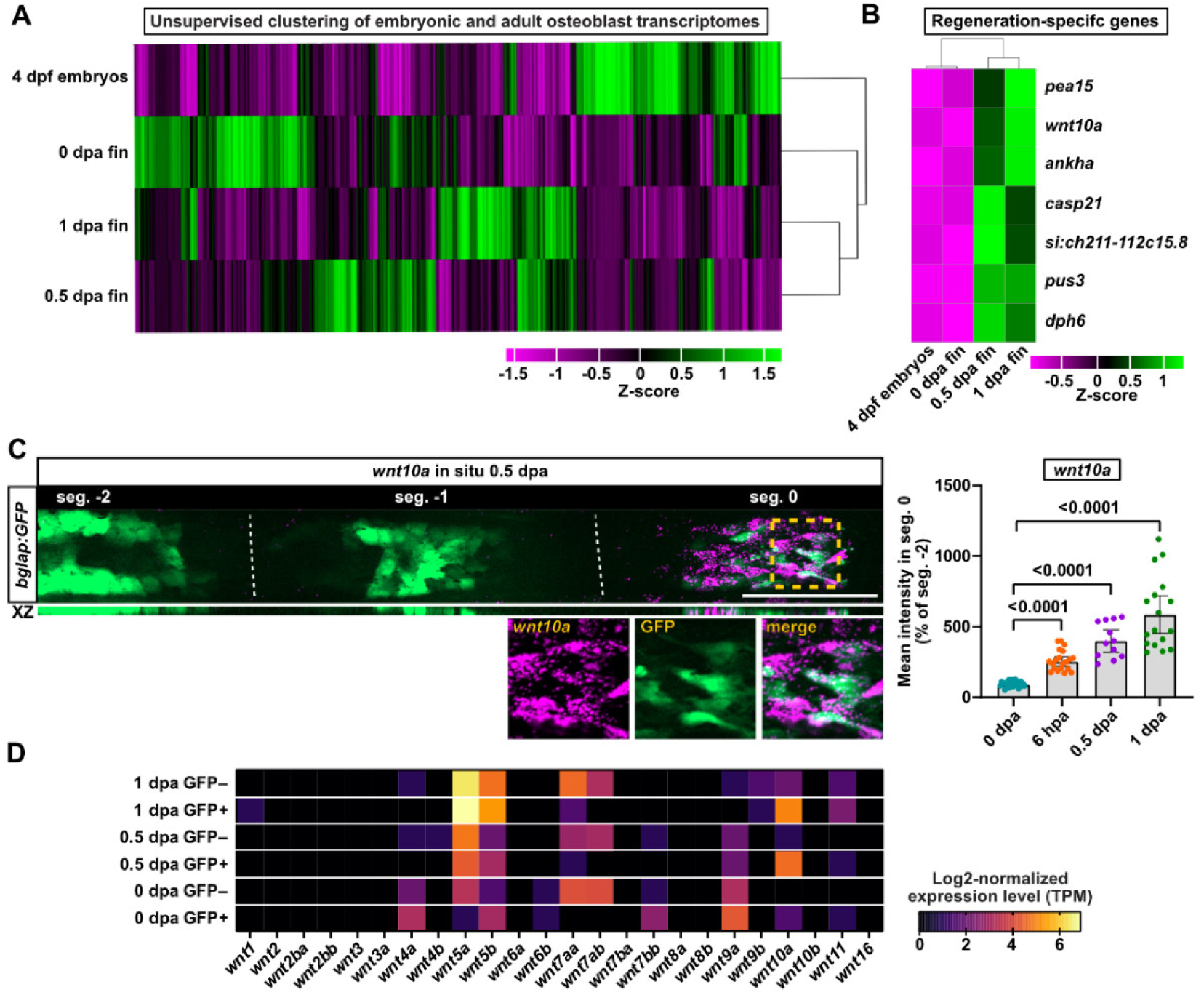
Osteoblasts dedifferentiate into a regenerative state that expresses *wnt10a*. **A)** Unsupervised clustering of z-score standardized gene expression of osteoblasts sorted from 4 days post fertilization (dpf) embryos ^23^ and our bulk RNAseq data from adult fin osteoblasts. **B)** Heatmap of selected genes that are specifically expressed in dedifferentiating adult fin osteoblasts. **C)** HCR in situ hybridization shows expression of *wnt10a* in segment 0 osteoblasts (identified by location in the XZ view and by co-expression with *bglap:GFP*) at 0.5 dpa. Graph shows quantification of HCR pixel intensities in segment 0 relative to segment -2 at the indicated time points. 0 dpa (nE = 3, nA = 15 total, nR (rays) = 30 total), 6 hpa (nE = 2, nA = 10 total, nR = 20 total), 0.5 dpa (nE = 1, nA = 6, nR = 12), 1 dpa (nE = 2, nA = 10 total, nR = 17 total). Dashed line, joints. Scale bar, 100 µm. Data are presented as mean values. Error bars, Mean ± 95% CI. Two tailed Student’s t-test. **D)** Heatmap of TPM-normalized RNASeq counts of all zebrafish Wnt ligands indicates that *wnt10a* is the most prominently upregulated Wnt/β-catenin signaling ligand at 0.5 and 1 dpa.

### wnt10a is upregulated in dedifferentiating osteoblasts

To identify drivers of osteoblast dedifferentiation, we focused on genes that are upregulated in 0.5 and 1 dpa fin osteoblasts relative to 0 dpa, but are not highly expressed in embryonic osteoblasts (**Figure 4B, Supplementary Fig. 6B**). While these included the potential regulators of inflammatory signaling (*il34, casp21, si:ch211-112c15.8* – a homolog of *Tnfrsf1a*), protein biosynthesis quality control (*dph6, pus3*), and (oxidative) stress response (*atad1b, smfn*) noted above, we focused our attention on the Wnt ligand *wnt10a. Wnt10a* expression was confirmed by qRT-PCR to be up in GFP+ osteoblasts at 0.5 dpa and 1 dpa (**Figure 2F, Supplementary Figure 3A**). Our scRNASeq data showed much higher expression of *wnt10a* in the “dedifferentiated osteoblasts” cluster than in differentiated osteoblasts and hardly any expression in any other cluster (**Figure 3G, Supplementary Figure 5B).** In situ hybridization confirmed that *wnt10a* is induced in osteoblasts in segment 0 by 6 hpa and that its expression increases by 0.5 dpa and 1 dpa (**Figure 4C**). *wnt10a* transcription has previously been recognized as being induced very early after fin amputation ^21^. Yet, the function of *wnt10a* during fin regeneration, and the role of Wnt/β-catenin signaling in early regenerative responses of cells of the stump to injury have not been investigated.

Analysis of our transcriptomics datasets revealed that of the 24 zebrafish Wnt ligands, only *wnt5a*, *wnt5b* and *wnt10a* were significantly upregulated in amputated fins at 0.5 and 1 dpa (**Figure 4D**). *wnt10a* was the only ligand specifically upregulated in osteoblasts (GFP+), and not in other cells (GFP–), while *wnt5a* and *wnt5b* were also strongly expressed in other cells (**Figure 4D**). Wnt5 ligands often activate β-catenin-independent pathways, which can counteract Wnt/β-catenin signaling ^24^, and *wnt5b* has been shown to attenuate fin regeneration via inhibition of the Wnt/β-catenin pathway ^21^. In contrast, Wnt10a typically acts through the canonical β-catenin pathway. The fact that *wnt10a* is the only canonical Wnt ligand upregulated early after fin amputation, combined with its osteoblast-specific expression prompted us to investigate the functional relevance of *wnt10a* and Wnt/β-catenin signaling in osteoblast dedifferentiation.

### Wnt/β-catenin signaling is necessary for osteoblast dedifferentiation

RT-qPCR for the direct Wnt/β-catenin target genes *lef1*, *axin2*, and *dkk1b* in *bglap:GFP*+ cells at 1 dpa indicated that Wnt/β-catenin signaling is active in dedifferentiating osteoblasts (**Supplementary Fig. 7A**). To inducibly interfere with Wnt/β-catenin signaling, we used a transgenic line in which mouse *axin1*, an inhibitor of the pathway, is expressed upon heatshock (*hsp70l:Mmu.Axin1-YFP^w^*^35^, abbreviated *hs:axin1*, **Figure 5A**) ^25^. RT-qPCR for the Wnt/β-catenin target genes *fgf20a*, *lef1*, and *axin2* confirmed inhibition of the pathway in fins at 1 dpa following three heat-shocks between -3 hpa and 21 hpa (**Figure 5B**). Interestingly, RT-qPCR also showed significantly higher expression of the osteoblast differentiation markers *bglapl*, *ccn4b and sparc* in *hs:axin1* transgenics compared to heat-shocked wild-type siblings, suggesting an impairment in osteoblast dedifferentiation (**Figure 5C**). This was confirmed by quantification of fluorescent in situ hybridization signals against the osteoblast differentiation markers *bglap, bglapl*, and *ccn4b* (**Figure 5D, Supplementary Fig. 7B**). Inhibition of Wnt/β-catenin impaired downregulation of differentiation markers both in the injured fin segment 0 and in the adjacent, non-injured segment -1 (**Figure 5D**).

**Figure 5.**
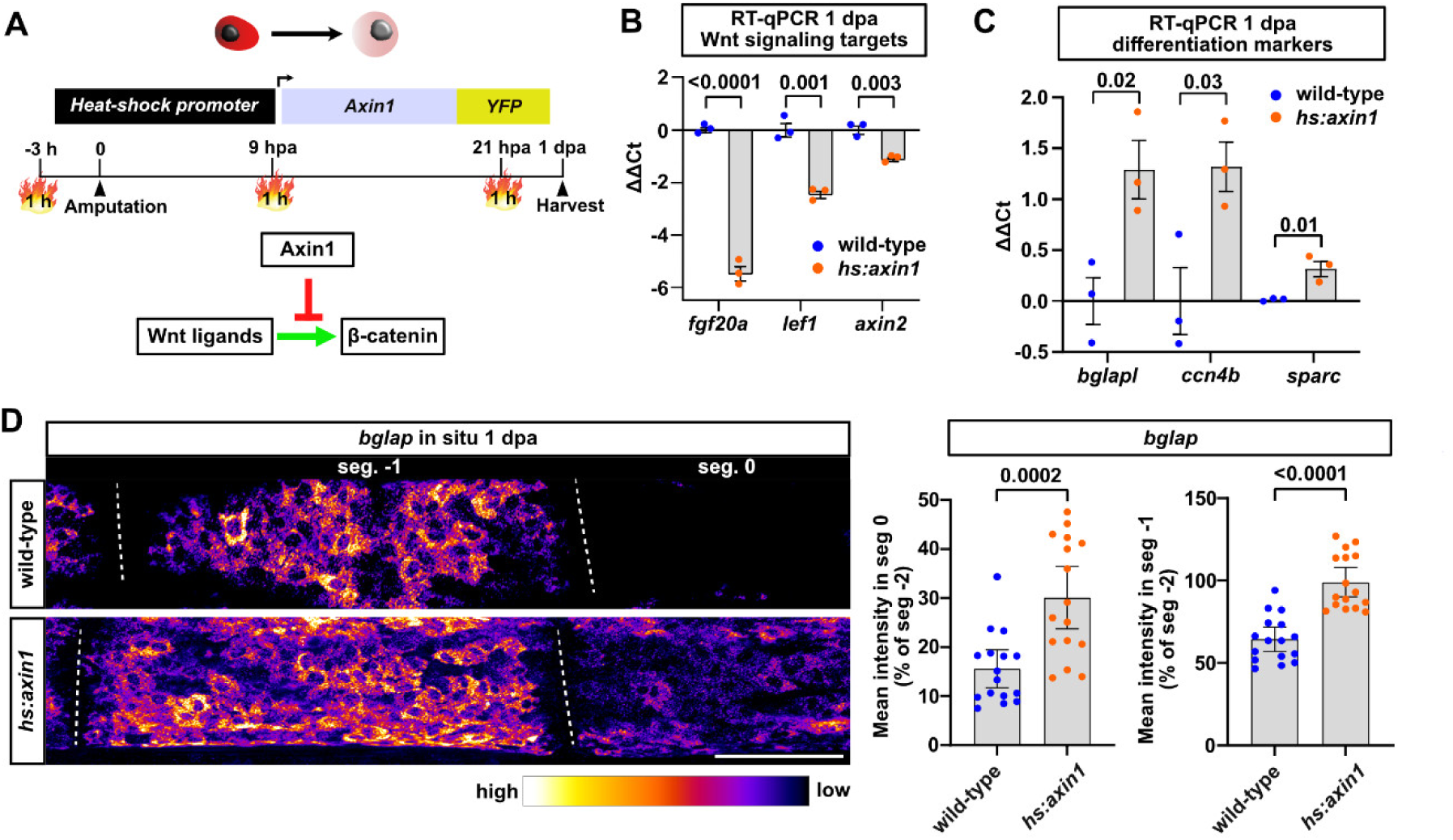
Inducible Wnt/β-catenin signaling loss-of-function impairs osteoblast dedifferentiation. **A)** *hs:axin1-YFP* fish were heat-shocked according to the scheme to induce Wnt/β-catenin signaling loss-of-function via expression of Axin1-YFP. **B)** RT-qPCR reveals downregulation of direct Wnt/β-catenin target genes in fin samples of *hs:axin1-YFP* fish compared to heat-shocked wildtypes at 1 dpa. nE (biological replicates) = 3, nA = 5 per replicate. **C)** RT-qPCR shows that osteoblast differentiation markers are more strongly expressed in fin samples of *hs:axin1-YFP* fish at 1 dpa, indicating reduced osteoblast dedifferentiation. nE = 3, nA = 5 per replicate. **D)** HCR in situ hybridization reveals reduced downregulation of *bglap* in segment 0 and segment -1 of *hs:axin1-YFP* fins at 1 dpa. nE = 2, nA = 10 total per group, nR = 16 total per group. Data are presented as mean values. Error bars, mean ± 95% CI. Two tailed Student’s t-test. Dashed line, joints. Scale bar, 100 µm. **(B, C)** Data are presented as mean values. ΔΔCt values are shown relative to the mean of the wildtype samples. Error bars, mean ± SEM. Two tailed Student’s t-test.

### wnt10a is required and sufficient for osteoblast dedifferentiation

Our Wnt ligand expression analysis suggested that *wnt10a* is the only ligand responsible for Wnt/β-catenin activation in dedifferentiating osteoblasts. Thus, we asked whether *wnt10a* is required for osteoblast dedifferentiation, using *wnt10a* mutants ^26^ (**Figure 6A**). While homozygous mutant fish are adult viable, they display severe fin development defects ^26^; thus, we analyzed heterozygous fish. Intriguingly, we found that the downregulation of *bglap* in segment -1 observed in wild-type fish (intensity in segment -1 at 62 ± 3% of that in segment -2) was completely absent in *wnt10a*^+/–^ siblings (segment -1 intensity at 97 ± 3% of that in segment -2, **Figure 6A**). We also performed bulk RNA-Seq of osteoblasts isolated from the dedifferentiation zone of *wnt10a*^+/–^ fins carrying the *bglap:GFP* transgene (**Figure 6B**). Mining these data revealed that *wnt10a*^+/–^ mutants showed reduced down- or upregulation of almost all of our validated high-confidence dedifferentiation-associated genes (**Figure 6B**). Furthermore, unsupervised clustering of their entire transcriptome showed that their transcriptional profile was clearly distinct from that of wild-type dedifferentiating osteoblasts (**Supplementary Fig. 8A**). We conclude that *wnt10a* is required for initiation of the transcriptional state transition occurring during osteoblast dedifferentiation. These defects in osteoblast dedifferentiation in *wnt10a*^+/–^ mutants were associated with reduced Wnt/β-catenin signaling in osteoblasts, as evidenced by derichment of genes annotated to function in the pathway and by reduced expression of the Wnt target genes *axin2*, *lef1* and *dkk1b* (**Supplementary Fig. 8B, C**). Gene set enrichment analysis also indicated that several signaling pathways, including Hh, Insulin, Notch, FGF, mTOR and FoxO are positively regulated by Wnt/β-catenin signaling in osteoblasts, since genes indicative of these pathways were downregulated in *wnt10a*^+/–^ osteoblasts (**Supplementary Fig. 9**).

**Figure 6.**
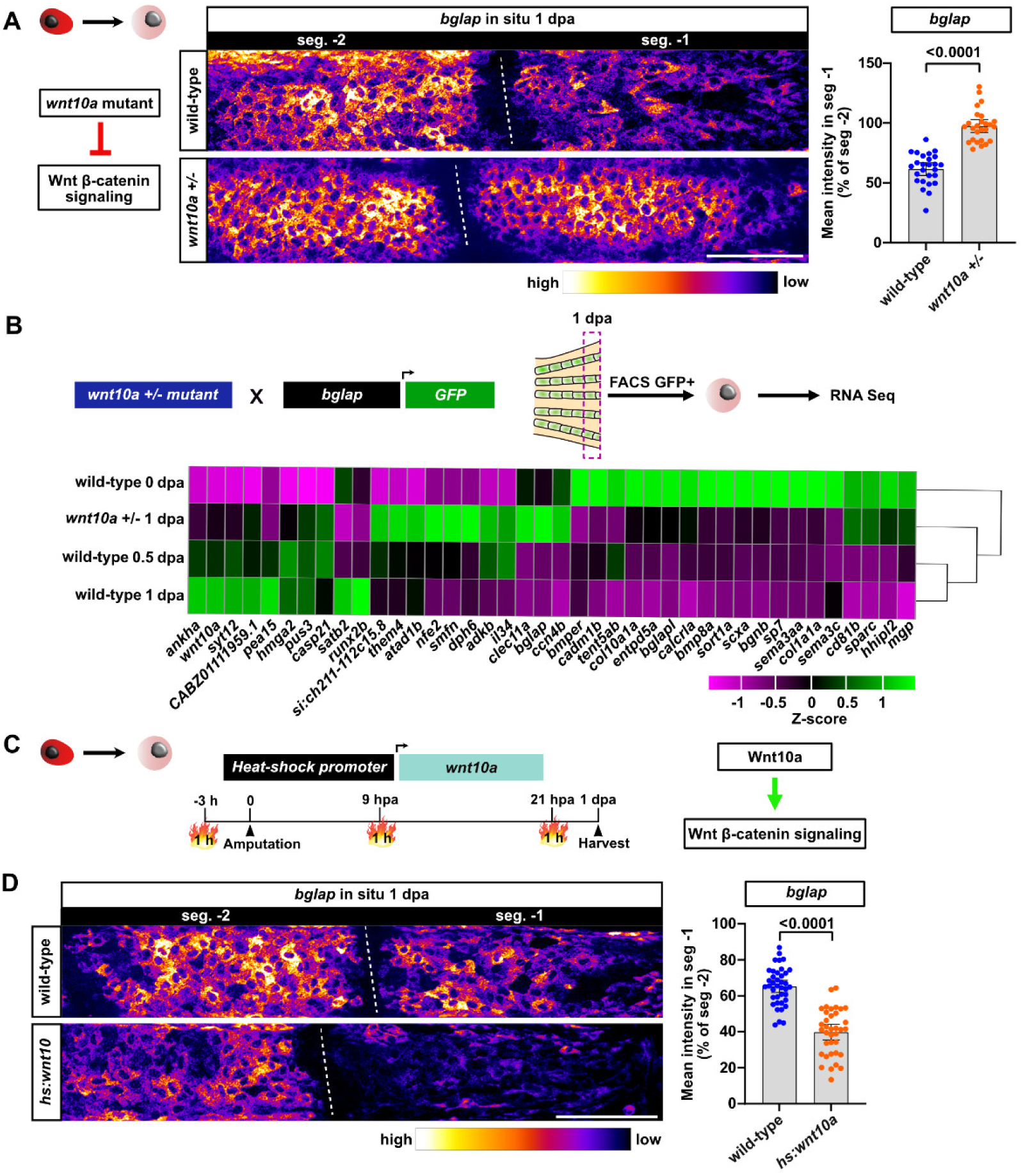
*wnt10a* mutants display osteoblast dedifferentiation defects, while wnt10a overexpression enhances dedifferentiation. **A)** HCR in situ hybridization shows reduced downregulation of *bglap* in segment -1 of *wnt10a^+/-^* mutants compared to wildtype siblings at 1 dpa. nE = 2, nA = 12 total per group, nR = 26 total per group. Dashed line, joints. Scale bar, 100µm. **B)** Unsupervised clustering of z-score standardized gene expression of osteoblasts sorted from *bglap:GFP; wnt10a*^+/-^ fish at 1 dpa with data from wild-type fish. Shown are expression values of those genes that we have validated by RT-qPCR to be down- or upregulated with dedifferentiation. **C)** *hs:wnt10a* fish were heat-shocked according to the scheme to induce Wnt/β-catenin signaling gain-of-function via expression of *wnt10a*. **D)** HCR in situ hybridization reveals increased downregulation of *bglap* in segment -1 of *hs:wnt10a* fins compared to heat-shocked wildtype siblings at 1 dpa. nE = 3, nA = 18 total per group, nR (wildtype) = 39 total, nR (*hs:wnt10a*) = 36 total. Dashed line, joints. Scale bar, 100 µm. **(A, D)** Data are presented as mean values. Error bars, mean ± 95% CI. Two tailed Student’s t-test.

Next, we asked whether *wnt10a* overexpression is sufficient to enhance osteoblast dedifferentiation, using the heat-shock inducible transgenic line *hsp70:wnt10a^stop^-EGFP ^fr60Tg^* (abbreviated *hs:wnt10a*). Three heat-shocks between -3 hpa and 21 hpa (**Figure 6C**) were sufficient to result in increased *wnt10a* expression in segment 0 and wide-spread ectopic expression in more proximal segments at 1 dpa (**Supplementary Fig. 10A**). Importantly, they also resulted in significantly stronger reduction of *bglap* expression in segment -1 (from 65 ± 2% of the level seen in segment -2 in heat-shocked wild-type siblings to 40 ± 2% in *hs:wnt10a* fins, **Figure 6D**). Furthermore, *wnt10a* overexpression was also sufficient to cause downregulation of *bglapl* in segment -1 at 1 dpa, whose expression in wild-types is only downregulated in segment 0 (**Supplementary Fig. 10B**). We conclude that overexpression of *wnt10a* is sufficient to enhance osteoblast dedifferentiation.

### Wnt/β-catenin signaling acts cell-autonomously to promote osteoblast dedifferentiation

Our results so far suggest that Wnt/β-catenin signaling acts cell autonomously in osteoblasts to promote their dedifferentiation. To test this, we made use of transgenic tools for cell-type specific conditional expression of *axin1*. First, we expressed mouse *axin1* in fin osteoblasts and fibroblasts (**Supplementary Fig. 11A, B**) after heat-shock, by combining a *her4.1(3.4):Cre, cryaa:YPet^ulm17Tg^* transgene (abbreviated *her4.1:Cre*) with a heat-shock inducible Cre responder line (*hsp70l:LOXP-Luciferase-MYC-STOP-LOXP-Mmu.-Axin1-p2A-NLS-dTomato,cryaa:AmCyan^ulm14Tg^*, abbreviated *hs:L to Axin1 nT* ^27^, **Figure 7A**). RT-qPCR showed that the Wnt target genes *fgf20a* and *axin2* were downregulated in heat-shocked double transgenics (**Figure 7B**). Importantly, quantification of *bglap* in situ signals showed that osteoblast dedifferentiation was completely inhibited in the double transgenics (**Figure 7C**). We conclude that Wnt/β-catenin signaling acts in fibroblasts and/or osteoblasts to regulate osteoblast dedifferentiation.

**Figure 7.**
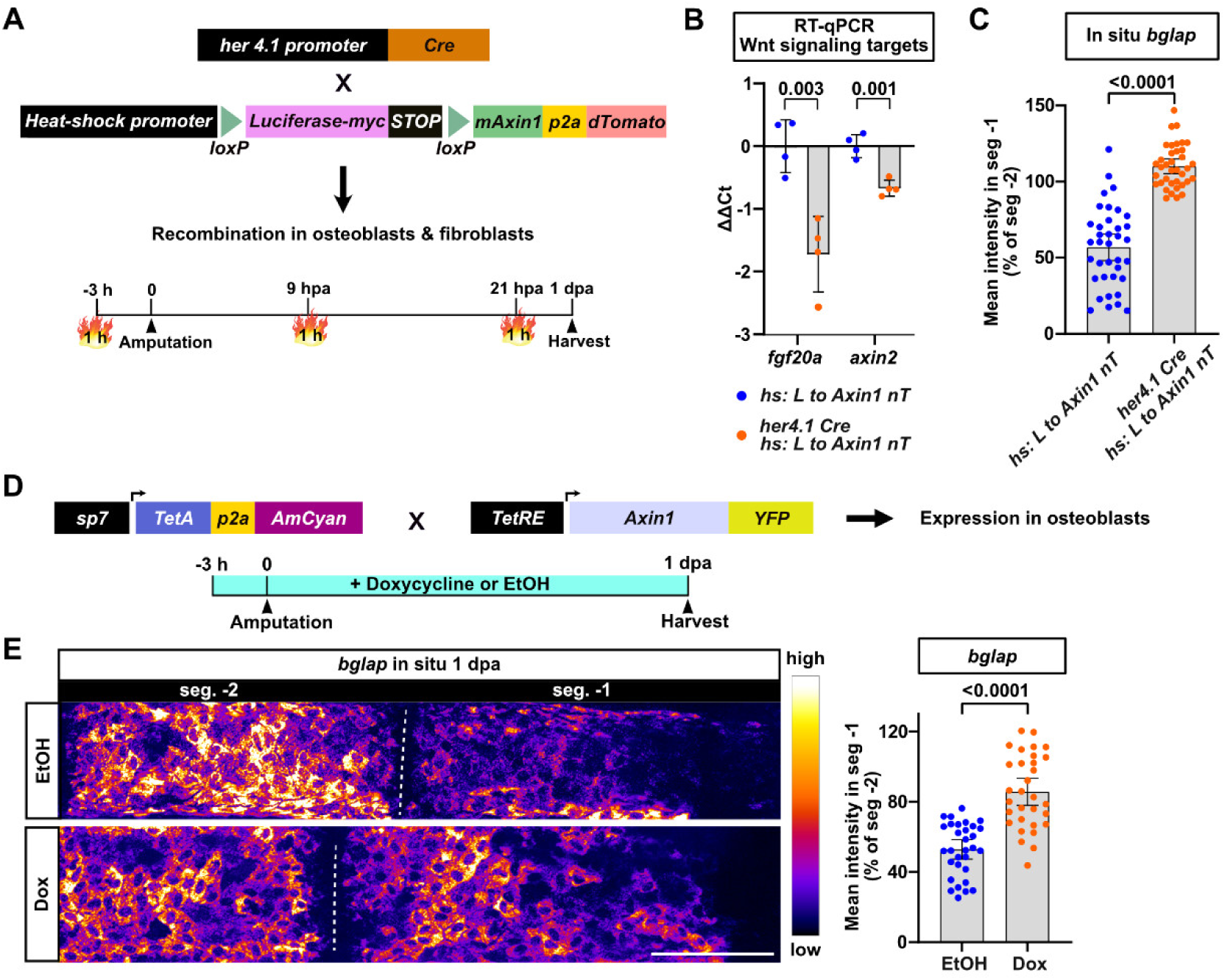
Wnt/β-catenin signaling is required in osteoblasts for their dedifferentiation. **A)** Cre-loxP transgenic strategy used to drive mouse Axin1 (mAxin1) and dTomato expression in fin osteoblasts and fibroblasts. **B)** RT-qPCR reveals downregulation of direct Wnt target genes in fin samples of heat-shocked double transgenic fish. nE (biological replicates) = 4, nA = 8 per replicate. ΔΔCt values are normalized to the mean of the single transgenic line. Error bars, mean ± SEM. Two tailed Student’s t-test. **C)** Quantification of HCR in situ hybridization signals reveals a complete lack of *bglap* downregulation in segment -1 of double transgenics. nE = 3, nA = 18 total per group, nR = 37 total per group. **D)** TetON system-based strategy used to inducibly express Axin1 in osteoblasts. **E)** HCR in situ hybridization reveals reduced downregulation of *bglap* in segment -1 of Doxycyclin (Dox) compared to EtOH treated double transgenic fish. nE = 3, nA = 18 total per group, nR = 31 total per group. Dashed line, joints. Scale bar, 100 µm. **(C, E)** Data are presented as mean values. Error bars, mean ± 95% CI. Two tailed Student’s t-test.

To specifically interfere with Wnt/β-catenin signaling in osteoblasts, we used the TetON system, combining the TetResponder line *TetRE:Mmu.Axin1-YFP^tud^*^1^ and a line expressing a TetActivator in osteoblasts (*sp7:irtTAM2(3F)-p2a-Cerulean^ulm3Tg^* ^19^, **Figure 7D**). While double transgenic fish treated with the vehicle control EtOH showed normal osteoblast dedifferentiation as quantified by *bglap* transcript downregulation in segment -1, induction of Axin1 expression by treatment with Doxycyclin (Dox) resulted in severe impairment of osteoblast dedifferentiation (**Figure 7E**). We conclude that Wnt/β-catenin acts cell-autonomously within osteoblasts to promote their dedifferentiation.

### Wnt/β-catenin signaling promotes osteoblast dedifferentiation via inhibition of NF-κB signaling

We have previously shown that NF-κB signaling acts cell-autonomously in osteoblasts to inhibit their dedifferentiation, and that the pathway needs to be downregulated for osteoblast dedifferentiation to occur ^14^. We thus wondered whether Wnt signaling promotes osteoblast dedifferentiation via inhibition of NF-κB signaling. Expression of the NF-κB signaling target genes *nfkbiaa* and *nfkbiab* was increased in osteoblasts sorted from *wnt10a*^+/–^ mutants at 1 dpa, as seen in our RNASeq data and verified by RT-qPCR (**Figure 8A**). Gene set enrichment confirmed that NF-κB signaling is downregulated in wild-type dedifferentiating osteoblasts at 1 dpa (**Supplementary Fig. 12A**), and showed that overexpression of *wnt10a* was able to further suppress NF-κB signaling in osteoblasts (**Supplementary Fig. 12B**). We conclude that Wnt/β-catenin signaling in osteoblasts inhibits NF-κB signaling. This implies that Wnt signaling acts upstream of NF-κB signaling in the regulation of osteoblast dedifferentiation.

**Figure 8.**
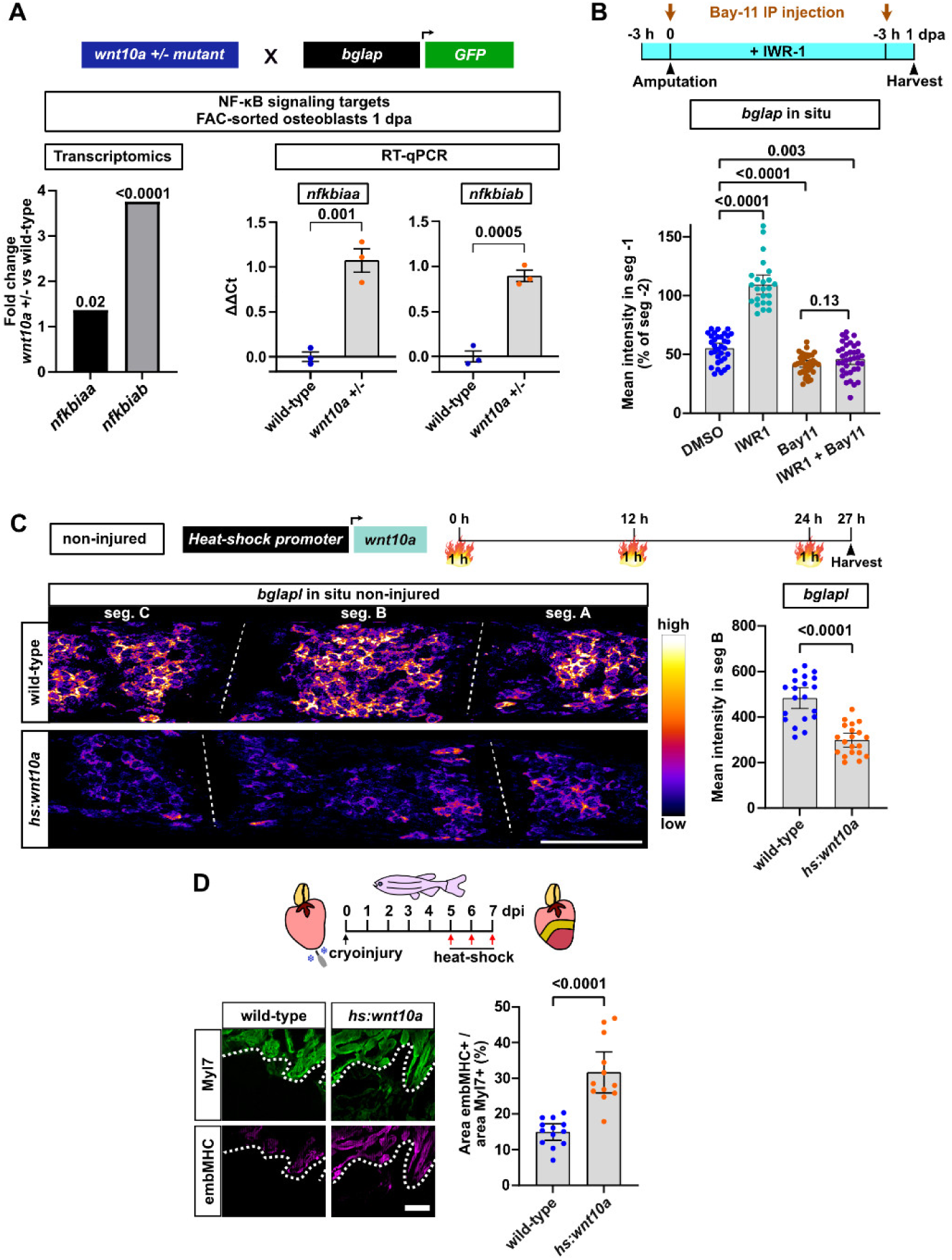
*wnt10a* induces osteoblast dedifferentiation via inhibition of NF-κB signaling, and is sufficient to induce dedifferentiation in non-injured fins and regenerating hearts. **A)** RNASeq (left) and RT-qPCR (right) reveal upregulation of the NF-κB signaling target genes *nfkbiaa* and *nfkbiab* in osteoblasts sorted from *bglap:GFP* fish heterozygous for the *wnt10a* mutation relative to their wild-type siblings at 1 dpa. nE (qPCR biological replicates) = 3, nA = 10 per replicate. ΔΔCt values are normalized to the mean of the wildtype at 1 dpa. Error bars, Mean ± SEM. Two tailed Student’s t-test. **B)** Osteoblast dedifferentiation, as measured by *bglap* downregulation in segment -1 revealed by HCR in situ hybridization, is inhibited in fish treated with the Wnt inhibitor IWR-1, while IP injection of the NF-κB inhibitor Bay-11 slightly but significantly enhances dedifferentiation. Treatment with both inhibitors yields results similar to those of Bay-11 alone. nE = 3 (except for 2 for IWR-1), nA = 18 total per group (12 for IWR-1), nR = 33 (DMSO), 24 (IWR-1), 37 (Bay-11), 33 (both). **C)** Overexpression of *wnt10a* using *hs:wnt10a* fish is sufficient to cause downregulation of *bglapl* detected by HCR in situ hybridization in non-injured fins, relative to heat-shocked wild-type fish. HCR signals were quantified in a bony segment (“B”) that is located at the same proximal-distal position as “segment -1” in amputated fins. nE = 2, nA = 12 total per group, nR = 22 total per group. Dashed line, joints. Scale bar, 100 µm. **D)** Immunofluorescence on cryosections of *hs:wnt10a* transgenic hearts reveals increased embryonic myosin heavy chain (embMHC) expression in Myl7+ cardiomyocytes at the wound border at 7 days post injury (dpi). Plots show the ventricular area covered by anti-embMHC staining relative to the 150 µm wound border zone area occupied by Myl7+ myocardium. nE = 2, nA = 13 wild-type, 11 *hs:wnt10a*. Scale bar, 100 µm. **(B, C, D)** Error bars, mean ± 95% CI. Two tailed Student’s t-test.

To test this, we used pharmacological interference with both pathways. We treated fish with a Wnt signaling inhibitor, IWR-1, or an NF-κB signaling inhibitor, Bay-11, or both, for one day and then performed in situ hybridization against *bglap* (**Figure 8B**). IWR-1 alone completely blocked osteoblast dedifferentiation (levels of *bglap* measured in segment -1 at 105 ± 5% of those detected in segment -2), as expected from the genetic Wnt loss-of-function experiments presented above (**Figure 8B**). In contrast, Bay-11 produced the opposite effect, further enhancing the already relatively strong dedifferentiation measured in DMSO-treated controls (from 56 ± 2% of the segment -2 levels to 41 ± 2%, **Figure 8B**), in accordance with our previously published findings ^14^. Interestingly, treating fish with both inhibitors produced the same effect as Bay-11 alone, enhancing dedifferentiation (*bglap* levels at 46 ± 2% of those in segment -2, **Figure 8B**). We conclude that Wnt/β-catenin signaling acts upstream of NF-κB signaling in the regulation of osteoblast dedifferentiation. Specifically, it appears to promote osteoblast dedifferentiation by inhibiting NF-κB signaling.

### Wnt10a expression is sufficient to induce osteoblast dedifferentiation in the absence of injury

Previously, we have shown that NF-κB signaling acts upstream of retinoic acid signaling in osteoblast dedifferentiation ^14^. Thus, the model we have built here implies that Wnt/β-catenin signaling acts at the top of the hierarchy of (known) regulators of osteoblast dedifferentiation. This prompted us to ask whether activation of Wnt/β-catenin signaling via *wnt10a* overexpression alone is sufficient to induce dedifferentiation without injury. We subjected non-injured *hs:wnt10a* fish and wild-type siblings to three heat-shocks within 24 h, harvested fins and compared *bglapl* levels between wild-type and transgenic fish (**Figure 8C**). Interestingly, we observed a severe downregulation of *bglap* expression in most bony segments of non-injured fins (**Figure 8C**). We conclude that *wnt10a* overexpression can induce osteoblast dedifferentiation in the absence of other injury-induced signals.

### Wnt10a promotes cardiomyocyte dedifferentiation

Dedifferentiation of mature cells provides source cells for regeneration also in several other highly regenerative systems, including the neonatal mouse and adult zebrafish hearts ^28,29^. In the regenerating heart, differentiated cardiomyocytes at the wound border lose characteristics of the differentiated phenotype, including organized sarcomeres, and upregulate embryonic genes such as embryonic myosin heavy chain (embMHC). We have previously shown that Wnt/β-catenin signaling is required for several aspects of cardiomyocyte regeneration, including their dedifferentiation ^27^. Given the ability of *wnt10a* to trigger osteoblast dedifferentiation, we asked whether it can also drive cardiomyocyte dedifferentiation. Intriguingly, compared to heat-shocked wild-type siblings, *hs:wnt10a* fish showed significantly increased cardiomyocyte dedifferentiation, as evidenced by the upregulation of embMHC in Myl7+ cardiomyocytes at the wound border at 7 day post injury (**Figure 8D**). Together these data show that *wnt10a* overexpression is sufficient to promote dedifferentiation of essential cell types that drive regeneration of the zebrafish heart and fin.

## Discussion

Dedifferentiation of mature cells into proliferative progenitors represents an intriguing and essential mechanism of source cell formation for regeneration of many systems. In this study, we establish sorting and bulk and single cell transcriptomics of dedifferentiating osteoblasts in regenerating zebrafish fins, which allowed us to derive the following insights. First, osteoblast dedifferentiation does not merely involve the downregulation of a few markers of differentiation and upregulation of some progenitor genes, but encompasses extensive transcriptional changes of thousands of genes. Second, osteoblasts do not dedifferentiate into a state that is identical to embryonic osteoblast progenitors. Rather, while they share many characteristics with such embryonic cells, they represent a regeneration-specific dedifferentiated state. One of the genes that we find to characterize this state is *wnt10a*. Using genetic and transgenic tools, we identify *Wnt10a*/β-catenin signaling as essential promoter of osteoblast dedifferentiation, that acts upstream of previously known regulators of this process. Specifically, our data suggest the following model for the regulation of osteoblast dedifferentiation (**Figure 9**).

**Figure 9.**
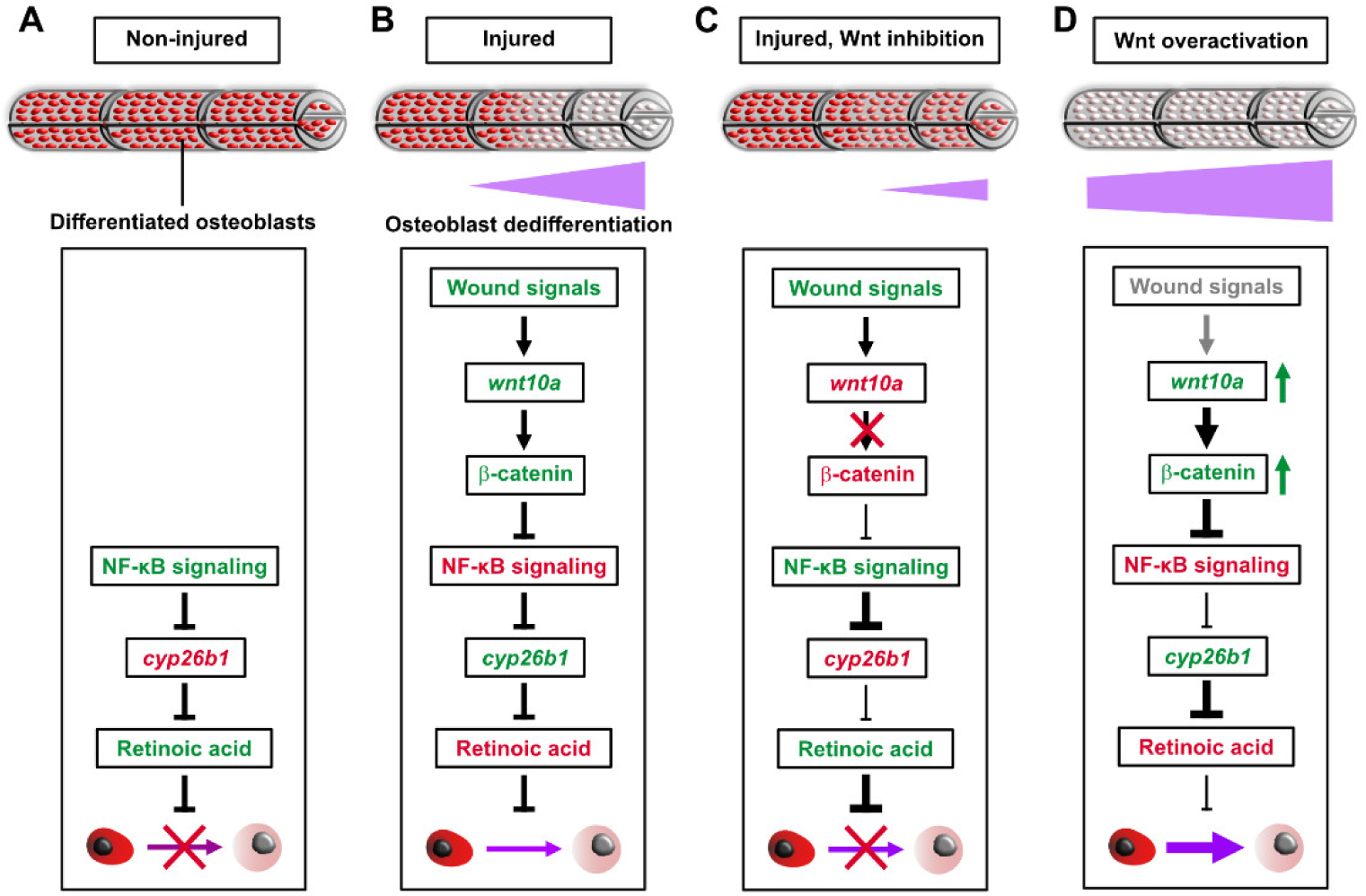
Model of *wnt10a* induced Wnt/β-catenin signaling function upstream of NF-κB signaling inhibition in osteoblast dedifferentiation. **A)** In non-injured fins, NF-κB signaling activity in differentiated osteoblasts suppresses expression of *cyp26b1*, allowing for retinoic acid signaling to be active, which maintains osteoblasts in the differentiated state ^14^. Green font indicates active pathway (components), red inactive ones. Thickness of lines indicates presumed strength of activating (arrows) or inhibitory interactions. **B)** Upon fin amputation, unidentified wound signals activate *wnt10a* transcription in osteoblasts. Through activation of the β-catenin signaling pathway, this results in inhibition of NF-κB signaling and subsequent graded dedifferentiation of osteoblasts in the injured and adjacent bony fin ray segment. **C)** Upon experimental reduction of *wnt10a* or inhibition of the β-catenin pathway, osteoblast dedifferentiation is blocked due to maintenance of NF-κB signaling activity. **D)** *Wnt10a* overexpression is sufficient to enhance dedifferentiation in injured fins and to induce it ectopically in the absence of wound signals in non-injured fins.

In the non-injured fin, NF-κB signaling is active in osteoblasts, where it prevents transcription of the retinoic-acid degrading enzyme *cyp26b1* ^14^. In the absence of *cyp26b1* expression, high levels of retinoic acid maintain osteoblasts in the differentiated state (**Figure 9A**) ^15^. In response to fin injury, *wnt10a* transcription is induced in osteoblasts located close to the amputation plane (“segment 0”) by unknown injury-induced cues. *wnt10a* activates the β-catenin signaling pathway, which acts cell-autonomously within osteoblasts to promote their dedifferentiation. It does so by inhibiting NF-κB signaling (**Figure 9B**). For reasons that we have not yet elucidated, dedifferentiation occurs in a graded fashion along the proximal-distal axis of the fin, reaching bone regions located hundreds of micrometers from the amputation plane and spreading into non-injured bone segments. When Wnt/β-catenin signaling is experimentally inhibited either by reducing *wnt10a* function or by osteoblast-specific overexpression of the pathway inhibitor Axin1, osteoblast dedifferentiation is blocked because NF-κB signaling activity increases (**Figure 9C**). Conversely, activation of the pathway via *wnt10a* overexpression increases osteoblast dedifferentiation by causing enhanced and ectopic NF-κB signaling inhibition (**Figure 9D**). Intriguingly, *wnt10a* overexpression is sufficient to trigger osteoblast dedifferentiation also in the absence of wound signaling in non-injured fins (**Figure 9D**). Thus, Wnt/β-catenin signaling, activated by *wnt10a,* triggers osteoblast dedifferentiation, acting upstream of a hierarchy of signals that maintain the differentiated state, whose loss induces dedifferentiation.

With this work, we identify a previously unknown essential role of Wnt signaling in one of the earliest regenerative responses of fins to injury that occurs in the stump of amputated fins. Of note, we and others have previously shown that Wnt/β-catenin signaling also regulates critical events that happen later during fin regeneration in the growing regenerate. In particular, it promotes proliferation of undifferentiated progenitor cells of the blastema and is also required for osteoblast re-differentiation ^19–21^. In both contexts – as regulator of osteoblast dedifferentiation in the stump and of later cellular functions in the regenerate – Wnt/β-catenin signaling acts upstream of several other signaling pathways. Intriguingly, however, in the regenerate Wnt/β-catenin signaling appears to largely act non-cell-autonomously; it sets up a signaling center that organizes regeneration via secretion of downstream signals ^19^. In contrast, we show here that the pathway regulates osteoblast dedifferentiation cell-autonomously. It even appears that the pathway is to a large extent activated in an autocrine manner by *wnt10a*, which is transcribed in the dedifferentiating osteoblasts themselves. Yet, our current evidence indicates that *wnt10a* is only transcribed in the injured segment 0, while Wnt/β-catenin-dependent osteoblast dedifferentiation occurs also in the adjacent segment -1. Currently, with existing experimental tools, we cannot distinguish between different potential explanations for this phenomenon. It is possible that we fail to detect weak transcription of *wnt10a* in segment -1 osteoblasts, where osteoblast dedifferentiation is not as complete as in segment 0 alternatively, Wnt10a protein might be transported to segment -1. While Wnt ligands typically cannot diffuse over extended distances because they carry hydrophobic modifications, several other mechanisms of Wnt ligand distribution have been discovered, including transport on cellular extensions ^30,31^. Although we show that inhibition of Wnt signaling in osteoblasts cell-autonomously interferes with osteoblast dedifferentiation, it is also possible that it directly acts only in the *wnt10a*-expressing osteoblasts in segment 0, and that its influence on osteoblasts in segment -1 is mediated by diffusible downstream signals, e.g. a diffusible inhibitor of NF-κB signaling.

Another interesting question raised by our work is to which extent the signals acting downstream of Wnt signaling in the early and late regenerative roles differ. While NF-κB signaling plays a prominent role in osteoblast dedifferentiation, a later role in the regenerate has not (yet) been described. On the other hand, retinoic acid signaling acts downstream of Wnt signaling in both contexts. Yet, in the regenerate, it mediates the positive effect of Wnt signaling on progenitor cell proliferation, and the retinoic acid producing enzyme *aldh1a2* is a direct transcriptional target of the pathway ^19^. Conversely, in the stump retinoic acid signaling appears to be indirectly inhibited by Wnt signaling via the NF-κB pathway and its regulation of *cyp26b1* transcription. Whether the other pathways that act downstream of Wnt signaling in the regenerate also play a role in osteoblast dedifferentiation (Activin, Notch, FGF, Hh, BMP)^19^ requires further investigation.

We propose that *wnt10a* is at the top of a hierarchy of signals known to regulate osteoblast dedifferentiation. This begs the question how *wnt10a* transcription is induced after fin amputation. We have previously shown that fin regeneration is triggered by the same (unidentified) generic injury-induced signals that also drive fin skin wound healing responses ^22^. Yet, osteoblast dedifferentiation versus osteoblast cell shape changes and migration towards the amputation plane are regulated independently, and a classical wound signal, activation of the complement system, is only required for osteoblast migration, but not their dedifferentiation ^6^. Of note, the *wnt10a* paralog *Wnt10b* has also been found to be induced by injury in mouse lung and muscle, and an enhancer partly mediating its injury responsiveness contains binding sites for the “immediate ealry” stress response AP-1 transcriptional complex ^32^. It will be interesting to study whether commonalities exist between injury-specific regulation of *wnt10a* and *wnt10b*, and to identify the molecular nature of the upstream signal(s).

Wnt/β-catenin signaling has emerged as essential regulator of regeneration in many systems, from hydra to mammals ^17,18,33^. While its exact functions in different systems vary, it has also previously been found to promote dedifferentiation of mature cells. For example, Wnt signaling promotes dedifferentiation of Müller glia cells during zebrafish retina regeneration ^34^. Also, Wnt/β-catenin signaling is required for cardiomyocyte dedifferentiation during zebrafish heart regeneration ^27^, and here we find that *wnt10a* overexpression is also sufficient to enhance cardiomyocyte dedifferentiation. Interestingly, Wnts have also been found to promote dedifferentiation of differentiated colorectal cancer cells to cancer stem cells, indicating that misregulated Wnt-mediated dedifferentiation can drive tumor formation ^35^.

An interesting question raised by the phenomenon of naturally occurring cellular dedifferentiation in regenerative processes is how “deep” cells dedifferentiate, that is whether they turn into an embryonic progenitor state or retain a partially differentiated character, and if they even dedifferentiate into a multipotent (embryonic) stem cell state. Alternatively, they might dedifferentiate into a state that resembles neither embryonic nor adult cells, but represents a special regenerative or repair state. Of note, in most cases of naturally occurring regenerative dedifferentiation, cells do not attain multipotency, but form source cells only for the cell type they formed from. This is the case for zebrafish and neonatal mouse cardiomyocytes during heart regeneration ^1,2,27^ newt muscle fibers during limb regeneration ^36^, and zebrafish osteoblasts during fin regeneration ^10^. Not surprisingly, there is little evidence that factors that are used to experimentally induce dedifferentiation and attainment of pluripotency of mature cells (iPSC formation) play a role in the known naturally occurring instances of cellular dedifferentiation. We also do not find any consistent upregulation of reprogramming factors in dedifferentiating osteoblasts, in fact expression of several such genes is downregulated with dedifferentiation (data not shown). Our comparison of the transcriptome of dedifferentiating adult and embryonic osteoblasts suggests that osteoblasts do not dedifferentiate into an embryonic state, although they gain many aspects of embryonic expression patterns. Rather, they appear to acquire a special regenerative state that is clearly distinct from both the adult differentiated and the embryonic state. Of note, this state includes strong expression (which we experimentally validated) of potential modulators of inflammatory signaling. This appears reminiscent of the repair state that Schwann cells dedifferentiate into following peripheral nerve injury in mammals, in which they secrete immune-modulatory cytokines and chemoattractants for immune cells ^37,38^. Possibly, osteoblasts do not only regulate their own dedifferentiation via production of *wnt10a*, but might play additional unappreciated roles in the regulation of regeneration, in particular in modulation of the immune response to injury.

In summary, our study identifies a novel role of *wnt10a*-driven Wnt/β-catenin signaling in the acquisition of regenerative properties by differentiated cells, and it suggests a potential role for dedifferentiated osteoblasts as instructive cells in the fin regeneration process.

## Methods

### Animals

All experiments involving zebrafish were approved by the state of Baden-Württemberg and the animal care representatives of Ulm University. Zebrafish were kept under standard conditions at 26-27°C water temperature and a 14/10 h light/dark cycle. If possible, mixed groups of males and females were used for all experiments. Since significantly different results between males and females were not observed in any of the experiments, data from both sexes are reported together. The following previously published transgenic and mutant fish lines were used: *Ola.Bglap:EGFP^hu^*^4008 10^, *hsp70l:Mmu.Axin1-YFP^w^*^35 25^, *hsp70:wnt10astop-eGFP^fr60Tg^*^26^, *hsp70l:LOXP-Luciferase-MYC-STOP-LOXP-Mmu.-Axin1-p2A-NLS-dTomato,cryaa:AmCyan^ulm14Tg^* ^27^, *TetRE:Mmu.Axin1-YFP^tud^*^1 19^, *sp7:irtTAM2(3F)-p2a-Cerulean^ulm3Tg^*^19^, and *wnt10a*^30922 26^. Transgenic or mutant fish were raised together with their wild-type siblings to adulthood; fish were genotyped and separated only shortly prior to any experiment, and wild-type siblings from the same cross were used as negative controls in all experiments.

### Creation of the her4.1 (3.4):Cre, cryaa:YPet^ulm17Tg^ transgenic line

The following DNA elements were assembled by PCR- and Gibson-assembly-based cloning methods: I-SceI meganuclease restriction site, MiniTol2 inverted repeat, zebrafish *cryaa* promoter, YPet, SV40 polyA signal, zebrafish *her4.1* promoter (3.4 kb) ^39^, zebrafish codon optimized Cre, SV40 polyA signal, MiniTol2 inverted repeat. Tol2 transposase-mediated transgene insertion was used to create a stable transgenic line. One subline was selected based on its ability to mediate recombination of several Cre responder lines in the embryonic central nervous system.

### Creation of the -1.5hsp70l:LOXP-Luciferase-MYC-STOP-LOXP-caegfra_L857R-2A-EGFP,cryaa:AmCyan^ulm20Tg^ transgenic line

A constitutively active zebrafish EGF receptor version (caEGFR) was created by exchanging L857 to R in zebrafish *egfra* (*erbb1*), ZDB-GENE-030918-1 using gene synthesis. The following DNA elements were assembled by PCR- and Gibson-assembly-based cloning methods: I-SceI meganuclease restriction site, MiniTol2 inverted repeat, attP site, *hsp70l* promoter (1.5kb), loxP, firefly luciferase 2, 6x myc tag, ocean pout antifreeze protein polyA signal, loxP, caEGFR, p2a, EGFP, SV40 pA signal, MiniTol2 inverted repeat. Tol2 transposase-mediated transgene insertion was used to create a stable transgenic line. One subline was selected based on its ability to be expressed in zebrafish osteoblasts after recombination driven by an osteoblast-specific Cre driver line, and by its ability to induce EGF signaling.

### Fin amputation

Adult zebrafish aged between 6 and 9 months were used for experiments. Caudal fin amputation was performed on fish anesthetized with 625 µM Ethyl 3-aminobenzoate methanesulfonate (Sigma-Aldrich, A5040) using sterile scalpels at approximately 50% fin length, which in most fins is just proximal to where fin rays bifurcate. Within the amputated segments, the amputation plane was chosen such that about 30-50% of the length of the amputated segment (“segment 0”) remained. This ensured that the osteoblast dedifferentiation zone encompassed approximately equal proportions of the non-amputated segment located proximally to segment 0 (“segment -1”) in all cases.

### Heart cryoinjury

Heart cryoinjuries were performed on fish 4–6 months of age that were anesthetized with 625 µM Ethyl 3-aminobenzoate methanesulfonate as previously described with a liquid nitrogen-cooled copper filament of 0.3-mm diameter ^40^. Injured fish were kept at a density of 7 fish per 1.5 liters.

### FACS sorting of osteoblasts

Differentiated osteoblasts were isolated at 1 day post amputation (dpa) from the whole stump or dedifferentiation zone (segment 0 and segment -1) of adult *bglap*:GFP transgenics using fluorescence-activated cell sorting (FACS) as described previously ^6^. Each of the three or four replicates for the bulk RNA sequencing experiments consisted of a pool of approximately 15 fins. In brief, samples were rinsed with cold Dulbecco’s phosphate buffered saline (DPBS) buffer (Thermo Fisher, 14190136), treated with 1× TrypLE Express enzyme (Thermo Fisher, 12604013) at 37° C for 30 min with gentle agitation, washed with cold DPBS buffer, and further dissociated into a single-cell suspension with 0.25 mg/mL Liberase DL (Merck, 5401160001) in enzyme-free Cell Dissociation Buffer (Thermo Fisher, 13151014) at 37° C for 30 min with gentle agitation. During this step, the samples were occasionally pipetted up and down using Pasteur pipettes with decreasing diameters to ensure complete digestion. The dissociated single-cell suspension was washed with cold eBioscience flowcytometry staining buffer (ThermoFisher, 00-4222-57) and pelleted by centrifugation at 500×g for 3 min at 4° C. Cells were resuspended in cold eBioscience flowcytometry staining buffer, and filtered through a 70 μm sample preparation filter (pluriSelect, 43-10070-46) to remove cell aggregates. The FACS Aria II flow cytometer was used to separate GFP+ and GFP− populations from single-cell suspensions. In order to define GFP background levels, wildtype non-transgenic fish were used. Forward and side scatter parameters were applied to exclude debris and doublets. Dead cells were excluded by staining with the cell-impermeable DNA-dye 4′,6-Diamidino-2-phenylindole dihydrochloride (DAPI, Sigma, 32670) and sorting for DAPI-negative cells. To further discriminate between GFP+ anuclear cell debris and intact cells, we stained with the cell-permeable DNA dye DRAQ5 (Cell Signaling, 4084S) and sorted for DRAQ5+ cells (gate shown in Supplementary Figure 1A). Therefore, DAPI−/DRAQ5+/GFP+ cells were sorted, and the same number of DAPI−/DRAQ5+/GFP− cells from each sample were also FAC-sorted to serve as control group.

For isolation and FAC-sorting of Zns5+ osteoblasts, segment 0 and -1 of caudal fins were harvested from approximately 35 wildtype fish per condition at 0 dpa, 6 hpa, 0.5 and 1 dpa, and digested according to the protocol described before. After resuspending and filtering the cells through a 70 μm sample preparation filter (pluriSelect, 43-10070-46), primary Zns5 antibody (ZIRC, ZDB-ATB-081002-37, RRID:AB_10013796) was added to the samples at a 1:100 dilution. The samples were then incubated at 4°C in the dark for 30 minutes. After incubation, they were centrifuged at 500 g for 5 minutes and washed twice with 2 ml of cold flow cytometry staining buffer followed by centrifugation at 500 g for 5 minutes. The cells were then resuspended in 100 µl flow cytometry staining buffer, and incubated with Goat anti-Mouse IgG (H+L) Highly Cross-Adsorbed Secondary Antibody (Invitrogen, A21424, RRID:AB_141780) at a 1:1000 dilution at 4°C for 30 minutes in the dark. Next, they were centrifuged at 500 g for 5 minutes, and washed twice with 2 ml of cold flow cytometry staining buffer. Cells were re-suspended in 300 μl of cold flow cytometry staining buffer. To distinguish live and dead cells, one drop of the viability dye, ReadiDrop 7-AAD (Bio-Rad, 1351102), was added to the cell suspension, followed by incubation for 2 minutes at room temperature in the dark. Cells gated by FACS as ZNS5+/7-AAD− were considered viable and were sorted into cold resuspension buffer; gate is shown in Supplementary Fig. 4A.

### RNA isolation

Total RNA was extracted from the FAC-sorted cells or fin segments using TRIzol reagent (Thermo Fisher, 15596018) according to the manufacturer’s protocol.

### Bulk RNA Sequencing cDNA synthesis

Low-input RNA cDNA synthesis and amplification was performed using the SmartSeq2 workflow. Total RNA was denatured for 3 min at 72° C in a total volume of 4 μL hypotonic buffer (0.2% Triton-X 100) with 2.4 mM dNTP (Invitrogen, 18427013), 240 nM dT-primer (5′AAGCAGTGGTATCAACGCAGAGTCGACT30VN-3′, where N represents a random base and V any base beside thymidine), and 4 U RNase Inhibitor (NEB, M0314). For reverse transcription, the reaction was supplemented with RT buffer, Template-Switch-oligo, and Superscript II reverse transcriptase, bringing the total volume to 10 µl with a final concentration of 1x Superscript II buffer (Invitrogen, 18064022), 1 M betaine, 5 mM DTT, 6 mM MgCl2, 1 µM TSO-primer (AAGCAGTGGTATCAACGCAGAGTACATrGrGrG, where rG stands for ribo-guanosine), 9 U RNase Inhibitor, and 90 U Superscript II. Reverse transcription was performed at 42° C for 90 min and subsequently inactivated at 70° C for 15 min. For subsequent PCR amplification of the cDNA, the optimal PCR cycle number was determined by qPCR using a 1 µl aliquot of the unpurified cDNA, along with 1X Kapa HiFi Hotstart Readymix (Roche, 07958927001), 1x SybrGreen, and 0.2 μM UP primer (AAGCAGTGGTATCAACGCAGAGT). Final amplification of residual cDNA was performed using 1X Kapa HiFi Hotstart Readymix under the following cycling conditions: initial denaturation at 98° C for 3 min, previously determined cycles (11-23 cycles) [98° C for 20 sec, 67° C for 15 sec, 72° C for 6 min], and final elongation at 72° C for 5 min. The amplified cDNA was purified using 0.6x volume of Sera-Mag SpeedBeads (GE Healthcare), resuspended in a buffer consisting of 10 mM Tris, 20 mM EDTA, 18.5% (w/v) PEG 8000, and a 2 M sodium chloride solution. The cDNA quality and concentration were assessed using the Fragment Analyzer NGS Fragment Kit (Agilent, DNF-473).

### Bulk RNA Sequencing library preparation

For library preparation, 2 µl of the amplified cDNA was tagmented with an equal volume of 1x Tagmentation Buffer and Bead-linked Transposome (Illumina DNA Prep, (M) Tagmentation, Illumina), adjusting with 0.4 µl of nuclease-free water for a total volume of 4 µl. The tagmentation reaction was conducted at 55° C for 15 min and subsequently stopped by incubating with 1 µl of 0.1% SDS at 37 °C for 15 min. DNA-binding beads were magnetically separated to discard the supernatant, and the bead-bound DNA was rehydrated in 15 µl of indexing PCR mix containing 1x KAPA Hifi HotStart Ready Mix and 300 nM of dual indexing primers (i5 and i7). Sample indexing PCR was performed under the following conditions: 72° C for 3 min, 98° C for 30 sec, followed by 12 cycles of [98° C for 10 sec, 63° C for 20 sec, 72° C for 1 min], and concluded at 72° C for 5 min. Libraries were purified at 0.9x ratios using Sera-Mag Speed Beads, followed by a 0.6x/0.9x size selection. Final libraries were quantified with the Fragment Analyzer NGS Fragment Kit (Agilent, DNF-473). Sequencing was performed on the Illumina NovaSeq 6000 using S4 flowcells in 100 bp paired-end XP mode, aiming for a minimum average sequencing depth of 30 million fragments per library.

### Bulk RNA Sequencing data analysis

Paired-end raw data (FASTQ files) were processed using HISAT2 in order to align the reads to the zebrafish genome assembly, GRCz11. To generate transcript-level counts Subread package featureCounts was used. Data normalization and differentially expressed gene analysis were performed using the R package DESeq2 ^41^. For Functional Enrichment Analysis, GO enrichment and Kyoto Encyclopedia of Genes and Genomes (KEGG) analysis were conducted using the R package clusterProfiler against the genome-wide annotation for Zebrafish (R package org.Dr.eg.db, DOI: 10.18129/B9.bioc.org.Dr.eg.db). To find interesting functional terms, the analysis was conducted using the “biological process” category of the GO, with the following parameters: an annotation cutoff of a minimum of 10 and a maximum of 500 counts, and a p-value cutoff of 0.05. The terms with FDR adjusted (P.adjust) value < 0.05 were considered as significantly enriched. The enrichment score plots were generated using the R package enrichplot (https://bioconductor.org/packages/release/bioc/html/enrichplot.html). For unsupervised clustering, DESeq2-normalized TPM counts were transformed to z-scores across samples and visualized as heatmaps using the R package pheatmap. Venn diagrams were generated based on log2 fold-change values using the criteria indicated in the corresponding figures.

### Single cell RNA sequencing

Chromium Next GEM Single Cell 3’ v3.1 kit (10x Genomics, 1000269) was used for cDNA synthesis and library preparations, according to the manufacturer’s instructions. In brief, 10000 cells from each condition were loaded into the single cell chip (10x Genomics, 1000127), and processed in the Chromium Controller. Downstream reactions were performed in Biorad C1000 Touch Thermal cyclers. 11 cycles were used for cDNA amplification. For cDNA clean-up, SPRIselect reagent (Beckman Coulter, B23317) was used. The cDNA quality and quantity were inspected using the Agilent Fragment Analyzer and the Agilent DNF-473 NGS Fragment Kit. 10 µl purified cDNA sample was used for library construction according to 10X Genomics protocol. Sample index PCR reaction was performed in Biorad C1000 Touch Thermal cyclers with 14 cycles. Libraries were evaluated using Agilent Fragment Analyzer and the Agilent DNF-473 NGS Fragment Kit. After quantification, the libraries were sequenced on the Illumina NovaSeq 6000 using S4 flow cells in 100 bp paired-end XP mode (R1/R2: 100 cycles; I1/I2: 10 cycles), generating an average sequencing depth of at least 280 million fragment pairs. Reads from all runs above, including 0 dpa, 6 hpa, 0.5 and 1 dpa were used in downstream analyses.

### Single cell RNA sequencing data analysis

Alignment of sequencing reads and processing into a digital gene expression matrix was performed using the 10x Genomics CellRanger version 7.2.0. The data was aligned against the zebrafish transcriptome GRCz11. Mapped reads for the 0 dpa, 6 hpa, 0.5 dpa and 1 dpa conditions were aggregated using the CellRanger pipeline prior to further computational analysis. Normalization and scaling of the count matrix were carried out using the R package Seurat v5.3.1 ^42^, utilizing the NormalizeData and ScaleData functions. Cells with either a low number of detected features (< 200 genes detected) or with abnormally high mitochondrial content (> 5%) were removed. The data were then normalized using a global-scaling normalization method, LogNormalize. Next, using the R package Seurat v5.3.1, highly variable features were selected, which were used for linear dimensional reduction analysis using Principal Component Analysis (PCA). The dimensionality of the dataset was determined using an elbow plot. The cells were then grouped into clusters using graph-based algorithms, and a non-linear dimensional reduction technique, UMAP, was run to visualize and explore the datasets. The clusters were then manually annotated by first identifying differentially expressed features and then finding the top cluster-specific marker genes. Feature plots for individual gene expression, dot plots of selected genes across all clusters as well as selected clusters, and heatmaps displaying the top 10 upregulated genes in each cluster were generated using the R package Seurat (v5.3.1).

### RT-qPCR

cDNA synthesis of RNA isolated from sorted cells or digested fin samples was conducted using the LunaScript RT SuperMIX Kit (New England Biolabs, E3010L) with random primers, after adjustment of RNA amounts to similar levels following quantification by UV absorbance using Nanodrop (Thermo Fisher Scientific). Quantitative real-time PCR was performed using the Luna Universal qPCR Master Mix (New England Biolabs, M 3003L) using primers listed in Supplementary Table 1. Ct values for each gene were normalized to the arithmetic mean of two housekeeping genes, *actin, beta 2* (ZDB-GENE-000329-3) and *hatn10* (ZDB-NUCMO-180807-7, and differential expression was calculated using the -2ΔΔCt method. Each biological sample was run in technical duplicates, and results were averaged. Primer amplification efficiency was determined on serial 1:4 dilutions of the cDNA mix prepared using the same tissue and conditions as those used for experimental samples.

### HCR whole mount in situ hybridization

Hybridization chain reaction (HCR) in situ hybridization was performed with the HCR RNA-FISH (v2.0) kit by Molecular Instruments and probes designed by and purchased from Molecular Instruments for the following genes: *bglap, bglapl, bgnb, ccn4b, clec11a, hmga2, nfe2, sparc,* and *wnt10a.* A table of identifiers used to design these probes can be found in Supplementary Table 2. Co-localization experiments were performed in *bglap:GFP* fish using the native GFP signal. To perform the HCR, fins were fixed overnight in 4% paraformaldehyde (PFA) at 4°C, washed several times with PBTw (1x PBS with 0.1% Tween), and pre-hybridized with 300 µL of probe hybridization buffer for 30 min at 37° C. This was replaced by probe solution by adding 2 µL of 1 µM probe stock solution to 300 µL of probe hybridization buffer at 37°C. The fins were incubated in the probe solution overnight at 37°C. After incubation, the fins were washed four times for 15 min each with 300 µL of probe wash buffer at 37°C, followed by two washes of 5 min each with 5x SSCT (SSC with 0.1% Tween) at room temperature. The samples were pre-amplified with 300 µL of amplification buffer for 30 min at room temperature. The hairpin solutions were prepared as follows: 30 pmol of hairpin h1 and 30 pmol of hairpin h2 were separately snap-cooled by heating 6 µL of 3 µM stock at 95°C for 90 seconds and cooling to room temperature for 30 min in the dark. The snap-cooled h1 hairpins and h2 hairpins were added to 300 µL amplification buffer at room temperature. The pre-amplification solution was removed and replaced with the hairpin solution. The fins were incubated overnight at room temperature in the dark. Excess hairpin solutions were removed by washing with 500 µL of 5× SSCT several times at room temperature. The fins were mounted on microscopic slides using FluorSave mounting reagent (Merck, 345789) and kept for at least 2 hours at room temperature before imaging.

### Heat-shock treatments

Details of the heat-shock regimes are provided in the figures and in the text of the relevant experiments. The heat-shocks were performed in automated tanks with heating from 27° C to at 37° C within 10 min, followed by incubation at 37° C for 1 h, after which water temperature was returned to 27° C within 15 min.

### Pharmacological interventions

Treatments of adult fish with the Wnt signaling inhibitor IWR1 (Merck, 681669) and with Doxycycline hyclate (Merck, D9891) were performed by incubation of fish in 1L of fish water. IWR-1 stock solution was prepared in DMSO (Merck, D2650) and used at 12 µM; control fish were treated with 0.1% DMSO. Doxycycline hyclate was dissolved in 50% EtOH and used at 1 µM; controls were treated with 0.05% EtOH. Bay-11 (Sigma-Aldrich, B5681) was applied by intraperitoneal injection. Bay-11 stock solution was prepared in DMSO, diluted to 50 µM in distilled water, of which 10 µl were injected; control fish were injected with 10 µl of 0.5% DMSO in distilled water.

### Heart immunostaining

Zebrafish hearts were fixed in 4% paraformaldehyde (PFA) (Merck, P6148) in phosphate buffer at RT for 1 h and processed as described in ^40^. Hearts were cryosectioned at a thickness of 10 μm. Sections were evenly distributed across six serial slides to ensure that each slide contained representative regions from all ventricular areas. For immunostaining, slides were washed three times with PEMTx buffer (80 mM Na-PIPES, 5 mM EGTA, 1 mM MgCl₂, pH 7.4, 0.2% Triton X-100) followed by a single wash with PEMTx containing 50 mM NH₄Cl. Blocking was performed in PEMTx supplemented with 10% Normal Goat Serum, 1% DMSO, and 89% PEMTx for 1 h at room temperature in a humidified chamber. Primary antibodies against embMHC Mouse monoclonal (MYH7 DSHB, N2.261, RRID:AB_531790), and Myl7 Rabbit polyclonal (GeneTex, GTX128346, RRID:AB_2885759) were diluted in PEMTx/normal goad serum and applied overnight at 4 °C. Secondary antibodies against Goat anti-Rabbit IgG (H+L) Cross-Adsorbed Secondary Antibody (Invitrogen, A21070, RRID:AB_2535731), and Goat anti-Mouse IgG (H+L) Highly Cross-Adsorbed Secondary Antibody (Invitrogen, A21424 RRID:AB_141780) were used at 1:1000. Nuclei were counterstained with DAPI (4′,6-diamidino-2-phenylindole). Slides were mounted with Fluorsave mounting medium (Merck, 345789).

### Imaging

Images were acquired using a Zeiss Axio Observer 7 microscope equipped with an Apotome using a 20x objective. High-resolution optical sections of whole-mount HCR-stained fins were collected to generate z-stacks, and all fin images shown are maximum intensity projections of the upper osteoblast layer of a single fin hemiray, while heart images were acquired as single optical planes.

### Image analysis

ImageJ Fiji was utilized for signal intensity quantification using the “mean gray value” function. For analysis of gene expression intensity along the proximodistal axis in HCR in situ images, five consecutive ROIs (segment height × 50 µm width) within segment 0 or segment -1 were analyzed with joints excluded, and signal intensity within these segments was normalized per ray to segment -2 of the same fin ray to account for imaging and processing variations. Quantification of the embMHC+ area on images of heart sections was restricted to the myocardium located within 150 µm of the wound border. The wound border was defined by cardiomyocyte marker immunofluorescence.

### Statistics

Statistical analyses were conducted using GraphPad Prism 9 software. Two-tailed Student’s t-tests were employed throughout. Exact p-values are reported in figures. A p-value < 0.05 was considered to indicate a statistically significant difference. Sample sizes are reported in figure legends as follows. nE = number of times the entire experiment was performed. In case of experiments where individual data points were derived by pooling of samples (e.g. RT-qPCR), nE reports the number of biological replicates. nA = sum total of animals used in all repeated experiments. In cases of experiments where data points were derived by pooling, nA reports the number of animals used per pool / replicate. nR = total number of fin rays analyzed.

## Data availability

Single cell and bulk RNASeq raw data are available at the gene expression omnibus (GEO) database, accession number GSE312516. Processed bulk RNASeq data, including the differentially expressed gene analyses results of different comparisons, are available in Supplementary Data File 1. Raw quantification data for all experiments can be found in the Supplementary “Source data” file. Newly created transgenic zebrafish lines adhere to nomenclature conventions set by the “Zebrafish Information Network, ZFIN”, and information about them can be found at ZFIN using the assigned allele numbers. Transgenic lines are available from the authors by request. Information on primary and secondary antibodies is compiled in Supplementary Tables 3 and 4.

## Acknowledgements

We thank Doris Weber, Janet Köhler, and Inna Guterman for their contributions to fish care, plasmid cloning, transgenic line establishment and maintenance. We would like to thank the ULMTeC Core Facility “Light microscopy” of the Medical Faculty at Ulm University for providing support and instrumentation for imaging funded by the Deutsche Forschungsgemeinschaft (DFG, German Research Foundation) – project number 257897648, and the core facility “cytometry” for support and instrumentation for cell sorting, funded by the DFG – project number 68236468. We thank Prof. Karin Danzer for granting us access to the Chromium Controller for 10x Genomics single-cell sequencing. We also acknowledge the DFG-funded NGS competence center DRESDEN-concept Genome Center (DcGC) for bulk and single cell RNASeq services. The Weidinger lab acknowledges funding by the DFG within the Collaborative Research Centre “Aging at Interfaces” (CRC1506; Project-ID 450627322), the CRC1149 “Danger Response, Disturbance Factors and Regenerative Potential After Acute Trauma” (Project-ID 251293561), and the CRC1279 “Exploiting the Human Peptidome for Novel Antimicrobial and Anticancer Agents” (project ID 316249678), and the project grant “Osteoblast heterogeneity in zebrafish regeneration”, project ID 51420450. H. F. M., D. P. P. and D.G. are or were members of the International Graduate School in Molecular Medicine, Ulm University.

## Contributions

Hossein Falah Mohammadi: Conceptualization (lead); Investigation – most experiments using zebrafish, Formal analysis – zebrafish transcriptomics data; Visualization; Writing – review and editing. Denise Posadas Pena: Heart experiments using *hs:wnt10a* in zebrafish. Dila Gülensoy: Visualization; Writing – review and editing. Ivonne Sehring: Validation of the *her4.1(3.4):Cre* zebrafish transgenic line. Writing – review and editing. Gilbert Weidinger: Conceptualization (lead); Funding Acquisition; Methodology – creation of *her4.1:Cre* and *hs:L 2 caEGFR-p2a-EGFP* zebrafish transgenic lines; Project Administration; Supervision; Visualization; Writing – original draft; Writing – review and editing.

## References

1 Porrello, E. R. et al. Transient regenerative potential of the neonatal mouse heart. Science 331, 1078–1080 (2011). 10.1126/science.1200708

2 Kikuchi, K. et al. Primary contribution to zebrafish heart regeneration by gata4(+) cardiomyocytes. Nature 464, 601–605 (2010). 10.1038/nature08804

3 Tanaka, H. V. et al. A developmentally regulated switch from stem cells to dedifferentiation for limb muscle regeneration in newts. Nature Communications 7, 11069 (2016). 10.1038/ncomms11069

4 Shivdasani, R. A., Clevers, H. & de Sauvage, F. J. Tissue regeneration: Reserve or reverse? Science 371, 784–786 (2021). 10.1126/science.abb6848

5 Merrell, A. J. & Stanger, B. Z. Adult cell plasticity in vivo: de-differentiation and transdifferentiation are back in style. Nat Rev Mol Cell Biol 17, 413–425 (2016). 10.1038/nrm.2016.24

6 Sehring, I. et al. Zebrafish fin regeneration involves generic and regeneration-specific osteoblast injury responses. Elife 11 (2022). 10.7554/eLife.77614

7 Geurtzen, K. et al. Mature osteoblasts dedifferentiate in response to traumatic bone injury in the zebrafish fin and skull. Development 141, 2225–2234 (2014). 10.1242/dev.105817

8 Stewart, S. & Stankunas, K. Limited dedifferentiation provides replacement tissue during zebrafish fin regeneration. Dev Biol 365, 339–349 (2012). 10.1016/j.ydbio.2012.02.031

9 Sousa, S. et al. Differentiated skeletal cells contribute to blastema formation during zebrafish fin regeneration. Development 138, 3897–3905 (2011). 10.1242/dev.064717

10 Knopf, F. et al. Bone regenerates via dedifferentiation of osteoblasts in the zebrafish fin. Dev Cell 20, 713–724 (2011). 10.1016/j.devcel.2011.04.014

11 Park, D. et al. Endogenous Bone Marrow MSCs Are Dynamic, Fate-Restricted Participants in Bone Maintenance and Regeneration. Cell Stem Cell 10, 259–272 (2012). 10.1016/j.stem.2012.02.003

12 Ando, K., Shibata, E., Hans, S., Brand, M. & Kawakami, A. Osteoblast Production by Reserved Progenitor Cells in Zebrafish Bone Regeneration and Maintenance. Developmental Cell 43, 643–650.e643 (2017). 10.1016/j.devcel.2017.10.015

13 Singh, Sumeet P., Holdway, Jennifer E. & Poss, Kenneth D. Regeneration of Amputated Zebrafish Fin Rays from De Novo Osteoblasts. Developmental Cell 22, 879–886 (2012). 10.1016/j.devcel.2012.03.006

14 Mishra, R., Sehring, I., Cederlund, M., Mulaw, M. & Weidinger, G. NF-κB Signaling Negatively Regulates Osteoblast Dedifferentiation during Zebrafish Bone Regeneration. Developmental Cell 52, 167–182.e167 (2020). 10.1016/j.devcel.2019.11.016

15 Blum, N. & Begemann, G. Retinoic acid signaling spatially restricts osteoblasts and controls ray-interray organization during zebrafish fin regeneration. Development 142, 2888–2893 (2015). 10.1242/dev.120212

16 Brandão, A. S. et al. A regeneration-triggered metabolic adaptation is necessary for cell identity transitions and cell cycle re-entry to support blastema formation and bone regeneration. eLife 11, e76987 (2022). 10.7554/eLife.76987

17 Walczyńska, K. S., Zhu, L. & Liang, Y. Insights into the role of the Wnt signaling pathway in the regeneration of animal model systems. Int J Dev Biol 67, 65–78 (2023). 10.1387/ijdb.220144yl

18 Petersen, C. P. Wnt signaling in whole-body regeneration. Curr Top Dev Biol 153, 347–380 (2023). 10.1016/bs.ctdb.2023.01.007

19 Wehner, D. et al. Wnt/β-catenin signaling defines organizing centers that orchestrate growth and differentiation of the regenerating zebrafish caudal fin. Cell Rep 6, 467–481 (2014). 10.1016/j.celrep.2013.12.036

20 Stewart, S., Gomez, A. W., Armstrong, B. E., Henner, A. & Stankunas, K. Sequential and opposing activities of Wnt and BMP coordinate zebrafish bone regeneration. Cell Rep 6, 482–498 (2014). 10.1016/j.celrep.2014.01.010

21 Stoick-Cooper, C. L. et al. Distinct Wnt signaling pathways have opposing roles in appendage regeneration. Development 134, 479–489 (2007). 10.1242/dev.001123

22 Owlarn, S. et al. Generic wound signals initiate regeneration in missing-tissue contexts. Nat Commun 8, 2282 (2017). 10.1038/s41467-017-02338-x

23 Raman, R. et al. The Osteoblast Transcriptome in Developing Zebrafish Reveals Key Roles for Extracellular Matrix Proteins Col10a1a and Fbln1 in Skeletal Development and Homeostasis. Biomolecules 14 (2024). 10.3390/biom14020139

24 Weidinger, G. & Moon, R. T. When Wnts antagonize Wnts. J Cell Biol 162, 753–755 (2003). 10.1083/jcb.200307181

25 Kagermeier-Schenk, B. et al. Waif1/5T4 Inhibits Wnt/β-Catenin Signaling and Activates Noncanonical Wnt Pathways by Modifying LRP6 Subcellular Localization. Developmental Cell 21, 1129–1143 (2011). 10.1016/j.devcel.2011.10.015

26 Benard, E. L. et al. wnt10a is required for zebrafish median fin fold maintenance and adult unpaired fin metamorphosis. Developmental Dynamics 253, 566–592 (2024). 10.1002/dvdy.672

27 Bertozzi, A., Wu, C.-C., Hans, S., Brand, M. & Weidinger, G. Wnt/β-catenin signaling acts cell-autonomously to promote cardiomyocyte regeneration in the zebrafish heart. Developmental Biology 481, 226–237 (2022). 10.1016/j.ydbio.2021.11.001

28 Jopling, C. et al. Zebrafish heart regeneration occurs by cardiomyocyte dedifferentiation and proliferation. Nature 464, 606–609 (2010). 10.1038/nature08899

29 O’Meara, C. C. et al. Transcriptional reversion of cardiac myocyte fate during mammalian cardiac regeneration. Circ Res 116, 804–815 (2015). 10.1161/circresaha.116.304269

30 Zhang, N. et al. Physiological regulation of neuronal Wnt activity is essential for TDP-43 localization and function. Embo j 43, 3388–3413 (2024). 10.1038/s44318-024-00156-8

31 Cooper, E. J. & Scholpp, S. Transport and gradient formation of Wnt and Fgf in the early zebrafish gastrula. Curr Top Dev Biol 157, 125–153 (2024). 10.1016/bs.ctdb.2023.12.003

32 Logan, C. Y. et al. Deletion of an enhancer that controls Wnt gene expression following tissue injury produces increased adipogenesis in regenerated muscle. Development 152 (2025). 10.1242/dev.204933

33 Narayanaswamy, S. & Technau, U. Self-organization of an organizer: Whole-body regeneration from reaggregated cells in cnidarians. Cells Dev 184, 204024 (2025). 10.1016/j.cdev.2025.204024

34 Ramachandran, R., Zhao, X. F. & Goldman, D. Ascl1a/Dkk/beta-catenin signaling pathway is necessary and glycogen synthase kinase-3beta inhibition is sufficient for zebrafish retina regeneration. Proc Natl Acad Sci U S A 108, 15858–15863 (2011). 10.1073/pnas.1107220108

35 Hu, Y. B. et al. Correction: Exosomal Wnt-induced dedifferentiation of colorectal cancer cells contributes to chemotherapy resistance. Oncogene 38, 6319–6321 (2019). 10.1038/s41388-019-0863-x

36 Sandoval-Guzmán, T. et al. Fundamental differences in dedifferentiation and stem cell recruitment during skeletal muscle regeneration in two salamander species. Cell Stem Cell 14, 174–187 (2014). 10.1016/j.stem.2013.11.007

37 Rutkowski, J. L. et al. Signals for proinflammatory cytokine secretion by human Schwann cells. J Neuroimmunol 101, 47–60 (1999). 10.1016/s0165-5728(99)00132-0

38 Tofaris, G. K., Patterson, P. H., Jessen, K. R. & Mirsky, R. Denervated Schwann cells attract macrophages by secretion of leukemia inhibitory factor (LIF) and monocyte chemoattractant protein-1 in a process regulated by interleukin-6 and LIF. J Neurosci 22, 6696–6703 (2002). 10.1523/jneurosci.22-15-06696.2002

39 Yeo, S. Y., Kim, M., Kim, H. S., Huh, T. L. & Chitnis, A. B. Fluorescent protein expression driven by her4 regulatory elements reveals the spatiotemporal pattern of Notch signaling in the nervous system of zebrafish embryos. Dev Biol 301, 555–567 (2007). 10.1016/j.ydbio.2006.10.020

40 Vasudevarao, M. D. et al. BMP signaling promotes zebrafish heart regeneration via alleviation of replication stress. Nature Communications 16, 1708 (2025). 10.1038/s41467-025-56993-6

41 Love, M. I., Huber, W. & Anders, S. Moderated estimation of fold change and dispersion for RNA-seq data with DESeq2. Genome Biology 15, 550 (2014). 10.1186/s13059-014-0550-8

42 Stuart, T. et al. Comprehensive Integration of Single-Cell Data. Cell 177, 1888–1902.e1821 (2019). 10.1016/j.cell.2019.05.031

