## Supplementary Figures and Tables for "Wnt/β-catenin signaling promotes zebrafish osteoblast dedifferentiation by *wnt10a*-mediated inhibition of NF-κB"

**Table of contents**

Supplementary Figure 1 page 2
Supplementary Figure 2 page 4
Supplementary Figure 3 page 5
Supplementary Figure 4 page 6
Supplementary Figure 5 page 7
Supplementary Figure 6 page 8
Supplementary Figure 7 page 9
Supplementary Figure 8 page 10
Supplementary Figure 9 page 11
Supplementary Figure 10 page 12
Supplementary Figure 11 page 13
Supplementary Figure 12 page 14

Supplementary Table 1 page 15
Supplementary Table 2 page 16
Supplementary Table 3 page 16
Supplementary Table 4 page 16

Supplementary References page 16

### Supplementary Figures

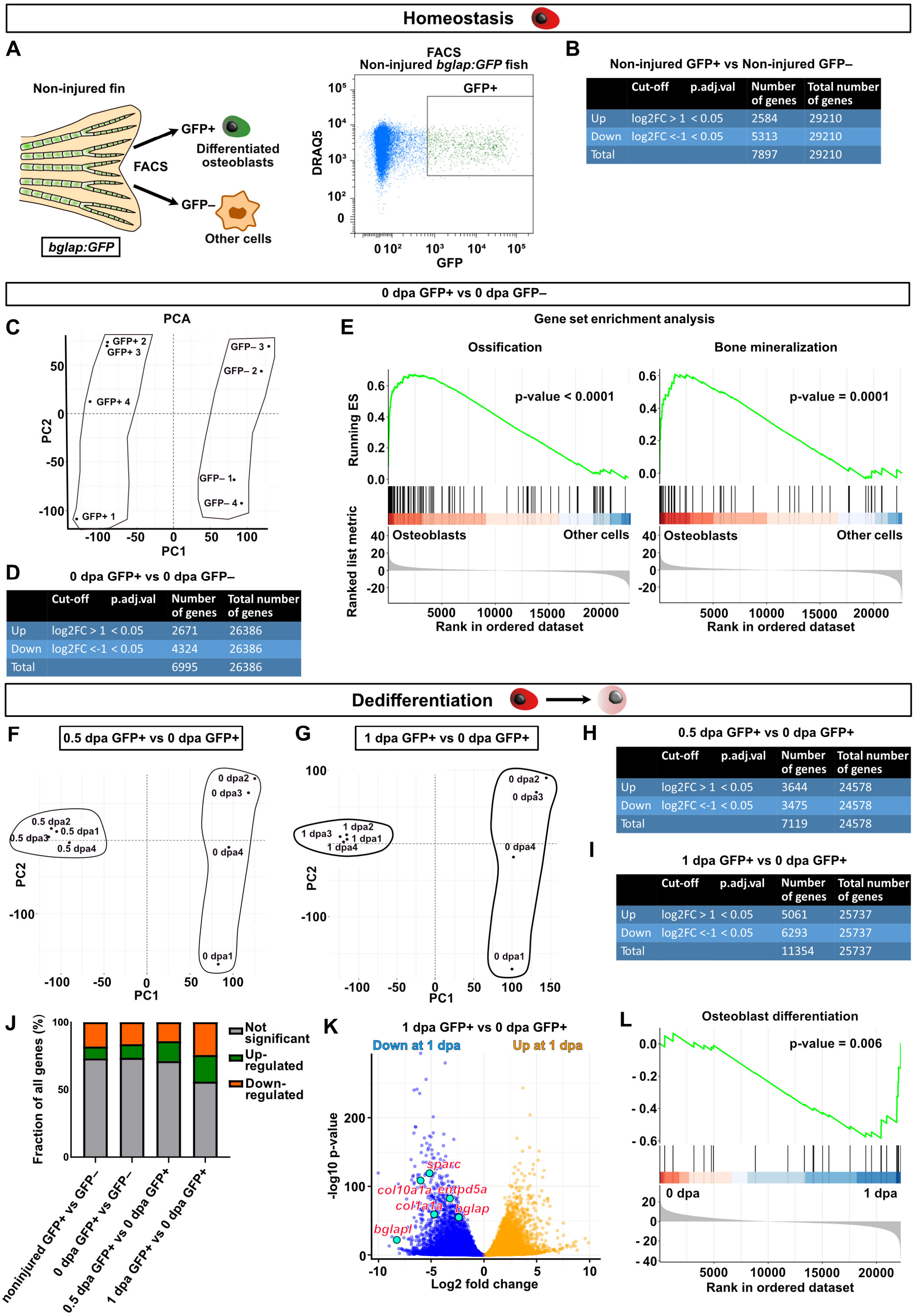


Non-injured GFP+ vs Non-injured GFP–

| | Cut-off | p.adj.val | Number of genes | Total number of genes |
|---|---|---|---|---|
| Up | log2FC > 1 | < 0.05 | 2584 | 29210 |
| Down | log2FC <-1 | < 0.05 | 5313 | 29210 |
| Total | | | 7897 | 29210 |

0 dpa GFP+ vs 0 dpa GFP–

| | Cut-off | p.adj.val | Number of genes | Total number of genes |
|---|---|---|---|---|
| Up | log2FC > 1 | < 0.05 | 2671 | 26386 |
| Down | log2FC <-1 | < 0.05 | 4324 | 26386 |
| Total | | | 6995 | 26386 |

0.5 dpa GFP+ vs 0 dpa GFP+

| | Cut-off | p.adj.val | Number of genes | Total number of genes |
|---|---|---|---|---|
| Up | log2FC > 1 | < 0.05 | 3644 | 24578 |
| Down | log2FC <-1 | < 0.05 | 3475 | 24578 |
| Total | | | 7119 | 24578 |

1 dpa GFP+ vs 0 dpa GFP+

| | Cut-off | p.adj.val | Number of genes | Total number of genes |
|---|---|---|---|---|
| Up | log2FC > 1 | < 0.05 | 5061 | 25737 |
| Down | log2FC <-1 | < 0.05 | 6293 | 25737 |
| Total | | | 11354 | 25737 |

#### Supplementary Figure 1.

**A)** FACS gating used to sort viable differentiated osteoblasts from entire non-injured adult caudal fins of *bglap*:GFP transgenic fish.

**B)** Numbers of genes found to be differentially expressed by the indicated filtering criteria between GFP+ and GFP– cells sorted from non-injured fins.

**C)** Principal component analysis graph showing that the transcriptomes of GFP+ and GFP– samples (quadruplicated) sorted from the dedifferentiation zone of *bglap:GFP* fins immediately after amputation (0 dpa) separate from each other on the PC1 axis, which explains most of the variations.

**D)** Numbers of genes found to be differentially expressed by the indicated filtering criteria between GFP+ and GFP– cells sorted from the dedifferentiation zone of injured *bglap:GFP* fins immediately after amputation (0 dpa).

**E)** Running enrichment score plots indicate upregulation of ossification and bone mineralization-related genes based on the GO terms "biological process" in GFP+ osteoblasts versus GFP– cells isolated at 0 dpa.

**F)** Principal component analysis graph reveals that the transcriptomes of GFP+ samples isolated at 0.5 dpa and 0 dpa are quite distant along the PC1 axis.

**G)** Principal component analysis graph reveals that the transcriptomes of GFP+ samples isolated at 1 dpa and 0 dpa are quite distant along the PC1 axis.

**H)** Numbers of genes found to be differentially expressed by the indicated filtering criteria between 0.5 dpa and 0 dpa GFP+ cells.

**I)** Numbers of genes found to be differentially expressed by the indicated filtering criteria between 1 dpa and 0 dpa GFP+ cells.

**J)** Number of differentially expressed genes relative to the total number of identifiable genes in the indicated comparisons.

**K)** Volcano plot illustrates downregulation of the known osteoblast mature marker genes in GFP+ osteoblasts at 1 dpa relative to 0 dpa.

L) Gene set running enrichment score plot indicating the downregulation of osteoblast differentiation-related genes based on the GO terms "biological process" at 1 dpa compared to 0 dpa.

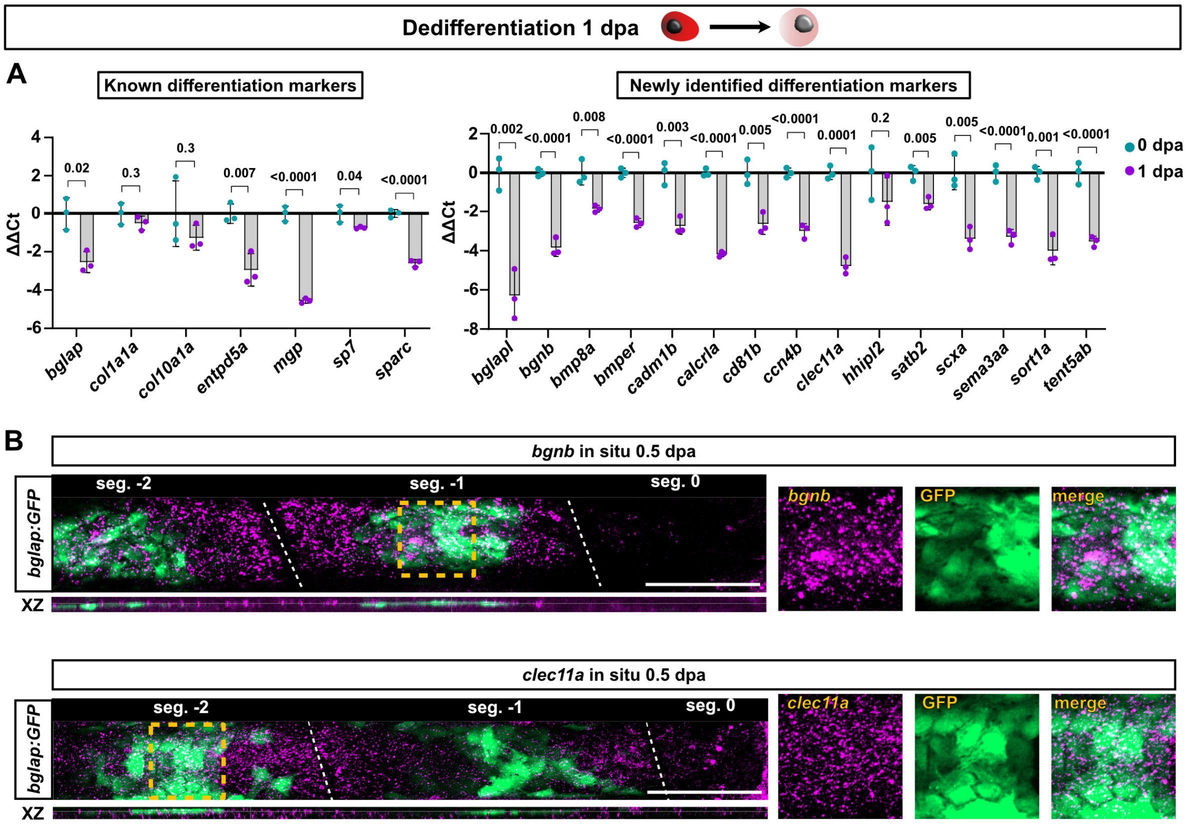


**Supplementary Figure 2.**

**A)** RT-qPCR confirms downregulation of known osteoblast differentiation genes (left graph), and newly identified genes that are specifically expressed in differentiated osteoblasts in non-injured fins (right graph) in GFP+ samples sorted from *bglap:GFP* fish at 1 dpa compared to 0 dpa. $n_E$ (biological replicates) = 3, $n_A$ = 15 per replicate. Data are presented as mean values. ΔΔCt values are normalized to the mean of the 0 dpa samples. Error bars, mean ± SEM. Two tailed Student´s t-test.

**B)** HCR in situ hybridization reveals expression of *bgnb* and *clec11a* in osteoblasts (identified by location in the XZ plot and by colocalization with *bglap:GFP*) at 0.5 dpa, and downregulation close to the amputation site. $n_E$ = 1, $n_A$ = 6, $n_R$ (rays) = 6. Dashed line, joints. Scale bar*,* 100 µm.

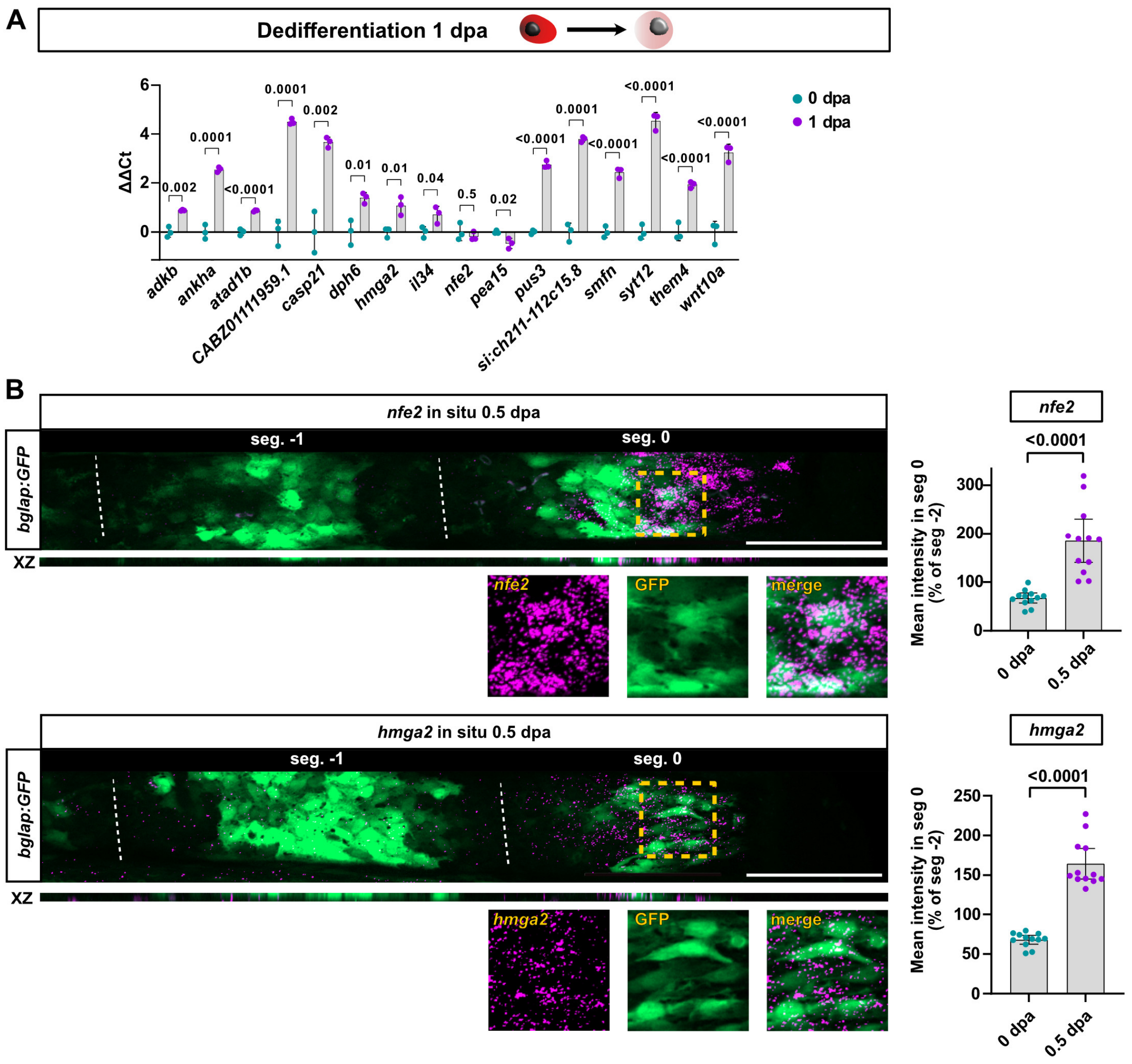


**Supplementary Figure 3.**

**A)** RT-qPCR confirms upregulation of genes identified by RNASeq in GFP+ dedifferentiating osteoblasts sorted from *bglap:GFP* fish at 1 dpa relative to GFP+ samples isolated at 0 dpa. $n_E = 3$ (biological replicates), $n_A = 15$ per replicate. Error bars, mean ± SEM. Two tailed Student´s t-test.

**B)** HCR in situ hybridization shows expression of *nfe2* and *hmga2* genes at 0.5 dpa in osteoblasts (identified by location in the XZ plot and by colocalization with *bglap:GFP*). Plots show quantification of HCR pixel intensities in segment 0 relative to segment -2 of the same rays. $n_E = 1$, $n_A = 6$, $n_R = 12$. Dashed line, joints. Scale bar, 100 µm. Data are presented as mean values. Error bars, mean ± 95% CI. Two tailed Student´s t-test.

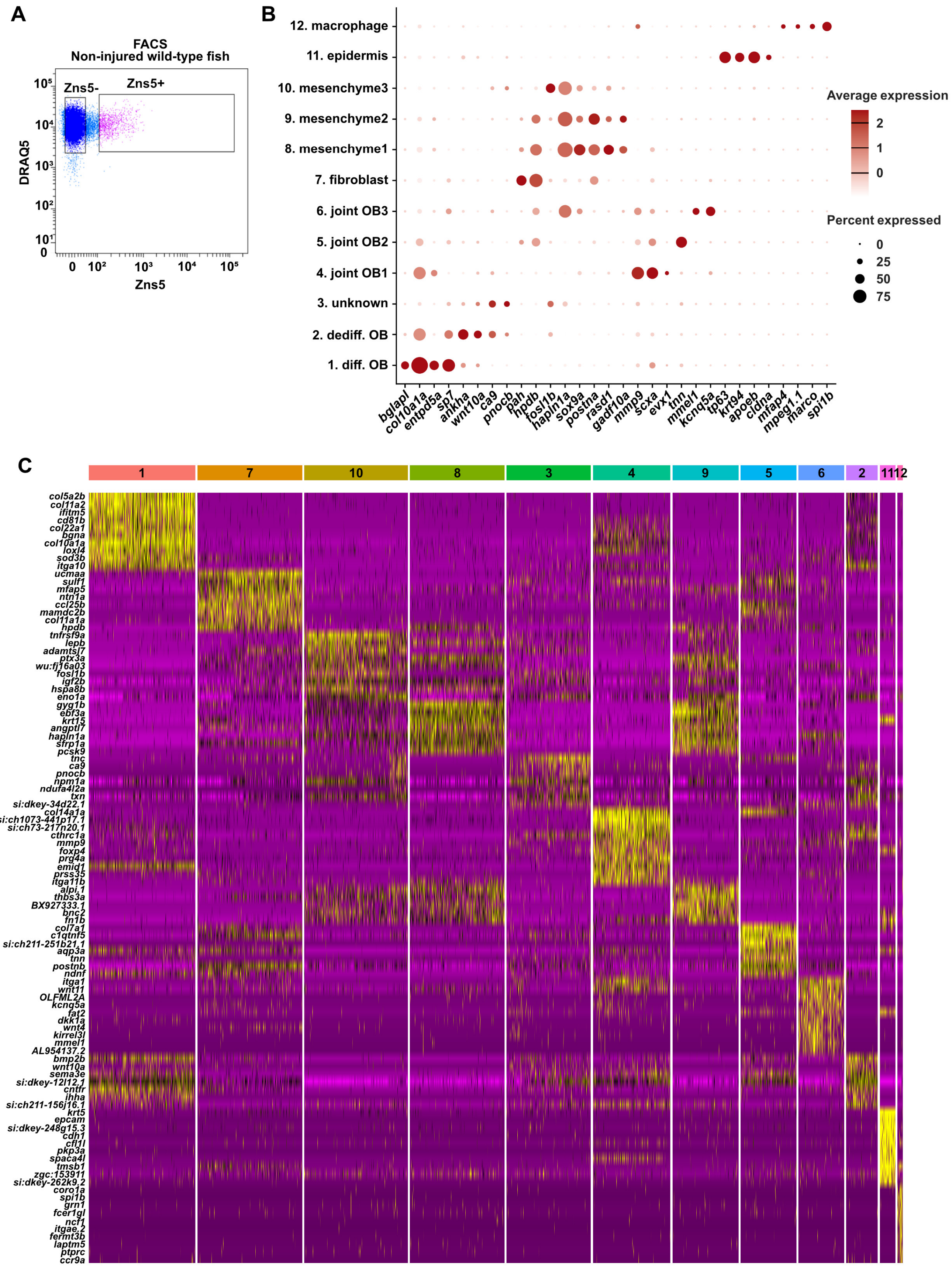


**Supplementary Figure 4.**

**A)** FACS gating used to sort viable Zns5+ and Zns5– cells from entire non-injured adult caudal fins.
**B)** Dot plot showing marker genes used for cluster annotation of single cell sequencing data.
**C)** Heatmap displaying the top 10 upregulated genes in each cluster.

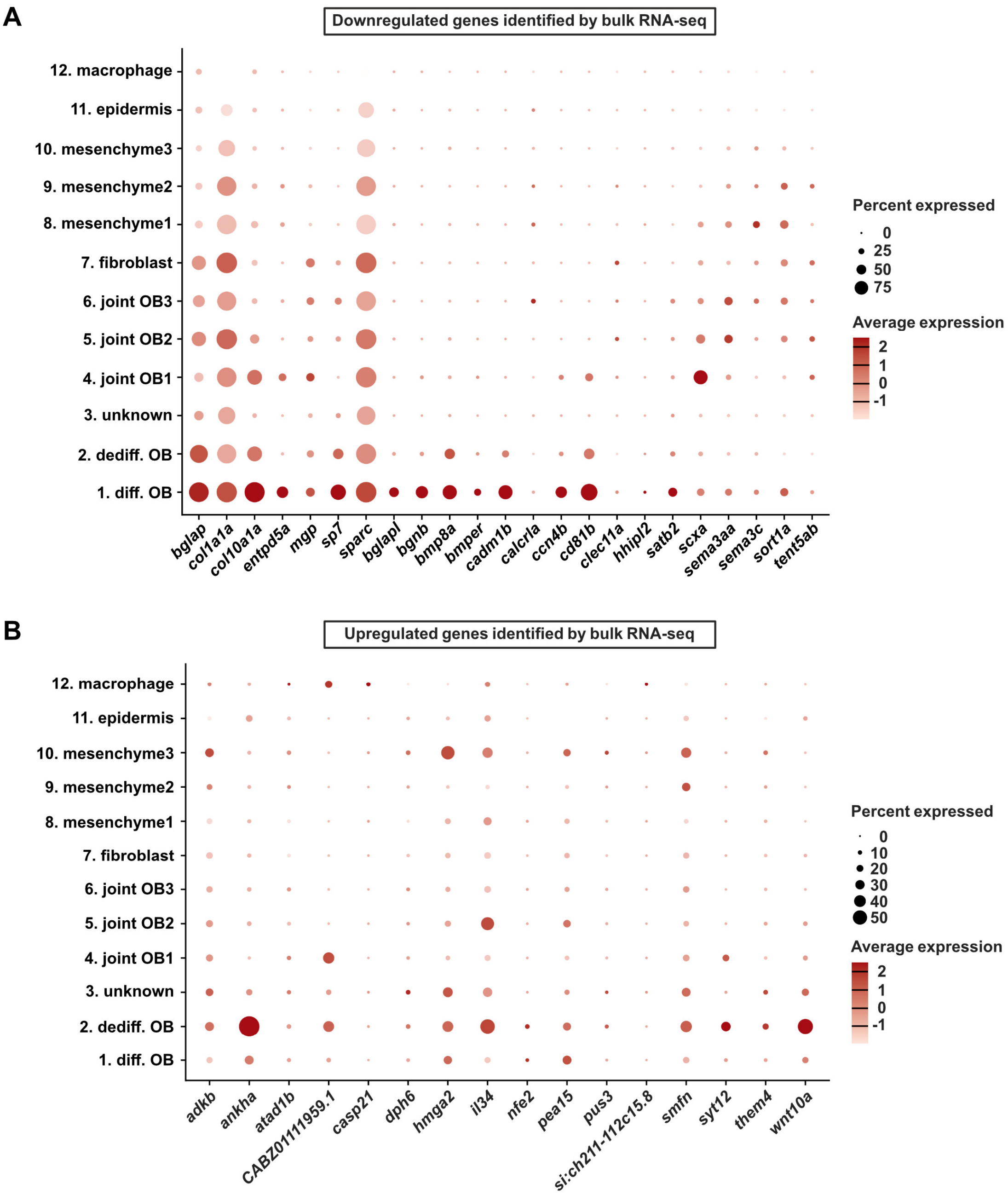


**Supplementary Figure 5.**

**A)** Dot plot comparing expression of genes that we have identified by bulk RNASeq to be downregulated in dedifferentiating osteoblasts in all of the clusters of the single cell data.

**B)** Dot plot comparing expression of genes that we have identified by bulk RNASeq to be upregulated in dedifferentiating osteoblasts in all of the clusters of the single cell data.

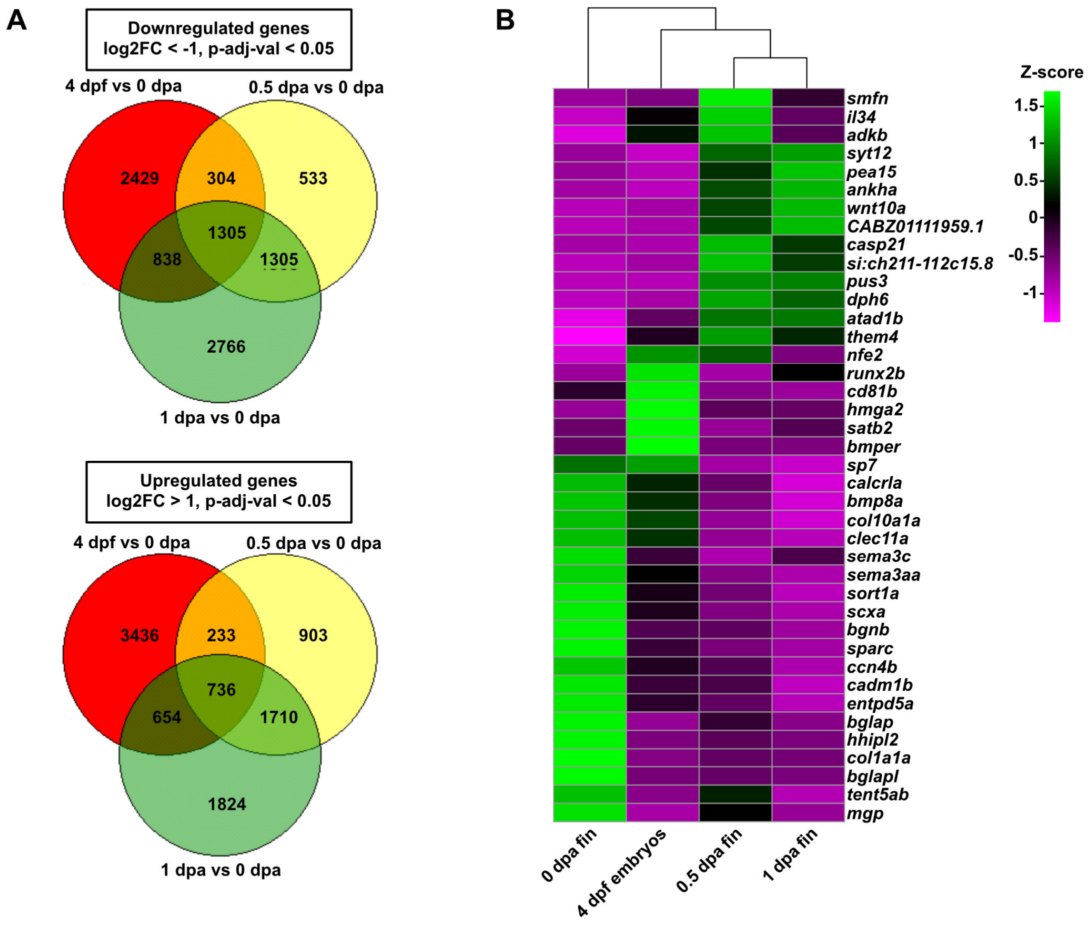


**Supplementary Figure 6.**

**A)** Venn diagrams showing overlap of genes found to be differentially expressed by the indicated criteria between the indicated bulk RNASeq datasets and 0 dpa *bglap:GFP+* adult fin osteoblasts. 0.5 dpa and 1 dpa represent our *bglap:GFP+* datasets, 4 dpf is a published sp7+ embryonic osteoblast dataset (Raman, et al., 2024).

**B)** Unsupervised clustering of the z-score standardized gene expression values of those genes that we have validated by RT-qPCR to be down- or upregulated in dedifferentiating adult fin osteoblasts in the indicated samples. Note that 0.5 and 1 dpa samples are more similar to the 4 dpf embryonic sample than to 0 dpa differentiated osteoblasts.

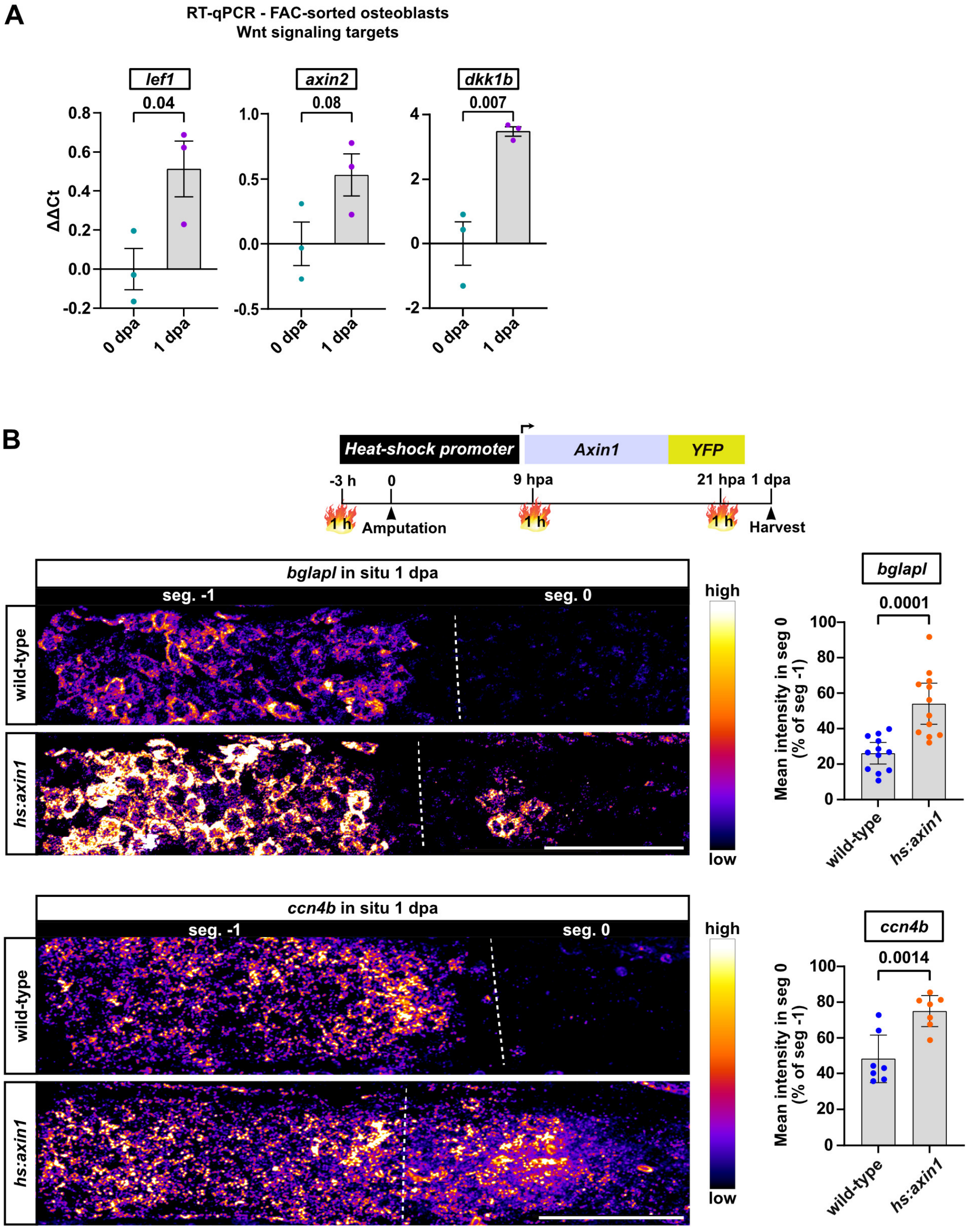


**Supplementary Figure 7.**

**A)** RT-qPCR indicates upregulation of direct Wnt/β-catenin target genes in sorted *bglap:GFP*+ osteoblasts at 1 dpa compared to 0 dpa. $n_E$ (biological replicates) = 3, $n_A$ = 15 per replicate. ΔΔCt values are normalized to the mean of the 0 dpa samples. Error bars, mean ± SEM. Two tailed Student´s t-test.

**B)** HCR in situ hybridization reveals reduced downregulation of the osteoblast differentiation markers *bglapl* and *ccn4b* in segment 0 of heat-shocked *hs:axin1-YFP* fish compared to heat-shocked wildtype siblings at 1 dpa, indicating reduced osteoblast dedifferentiation upon Wnt/β-catenin signaling inhibition. In wild-type fins, expression of *bglapl* and *ccn4b* is not downregulated in segment -1, which allowed us to use the intensities in segment -1 as reference for normalization of segment 0 signal intensities. $n_E$ = 1, $n_A$ = 6 per gene, $n_R$ = 12 (*bglapl*), 7 (*ccn4b*). Dashed line, joints. Scale bar, 100 µm. Data are presented as mean values. Error bars, mean ± 95% CI. Two tailed Student´s t-test.

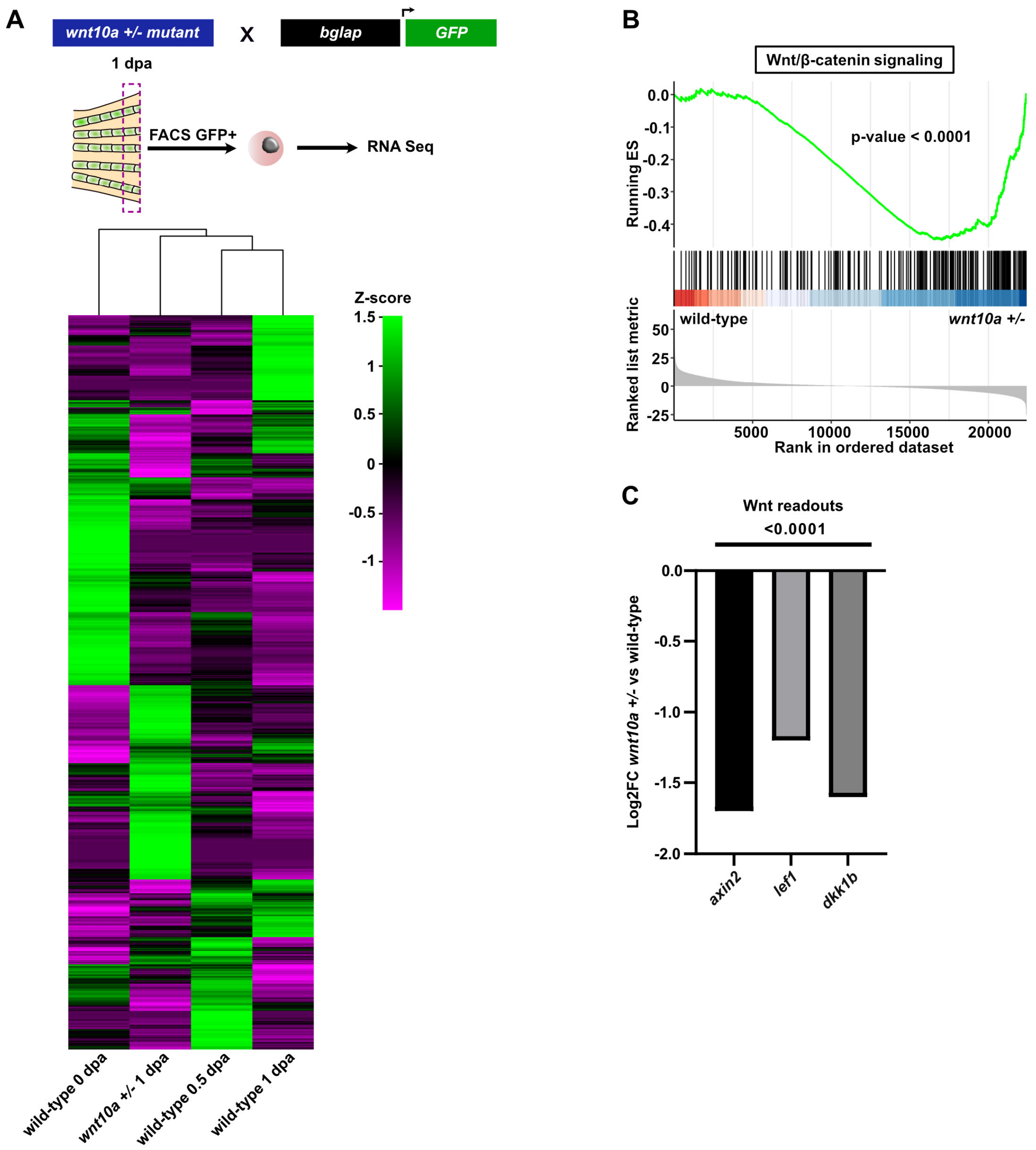


**Supplementary Figure 8.**

**A)** Unsupervised clustering of the z-score standardized gene expression values of all identifiable genes in our bulk RNASeq samples from 0, 0.5 and 1 dpa *bglap:GFP+* fin osteoblasts with another set derived from osteoblasts sorted from *bglap:GFP+; wnt10a* +/- fins at 1 dpa. Note that the *wnt10a*$^{+/-}$ sample exhibits a distinct expression profile positioned between 0 dpa and the 0.5 – 1 dpa wild-type samples.

**B)** Running enrichment score plot shows negative enrichment of genes annotated to function in Wnt/β-catenin signaling in 1 dpa osteoblasts of the *wnt10a* mutants.

**C)** Expression values of direct Wnt target genes are reduced in the RNAseq data of *wnt10a* mutant osteoblasts isolated at 1 dpa, relative to 1 dpa wild-type.

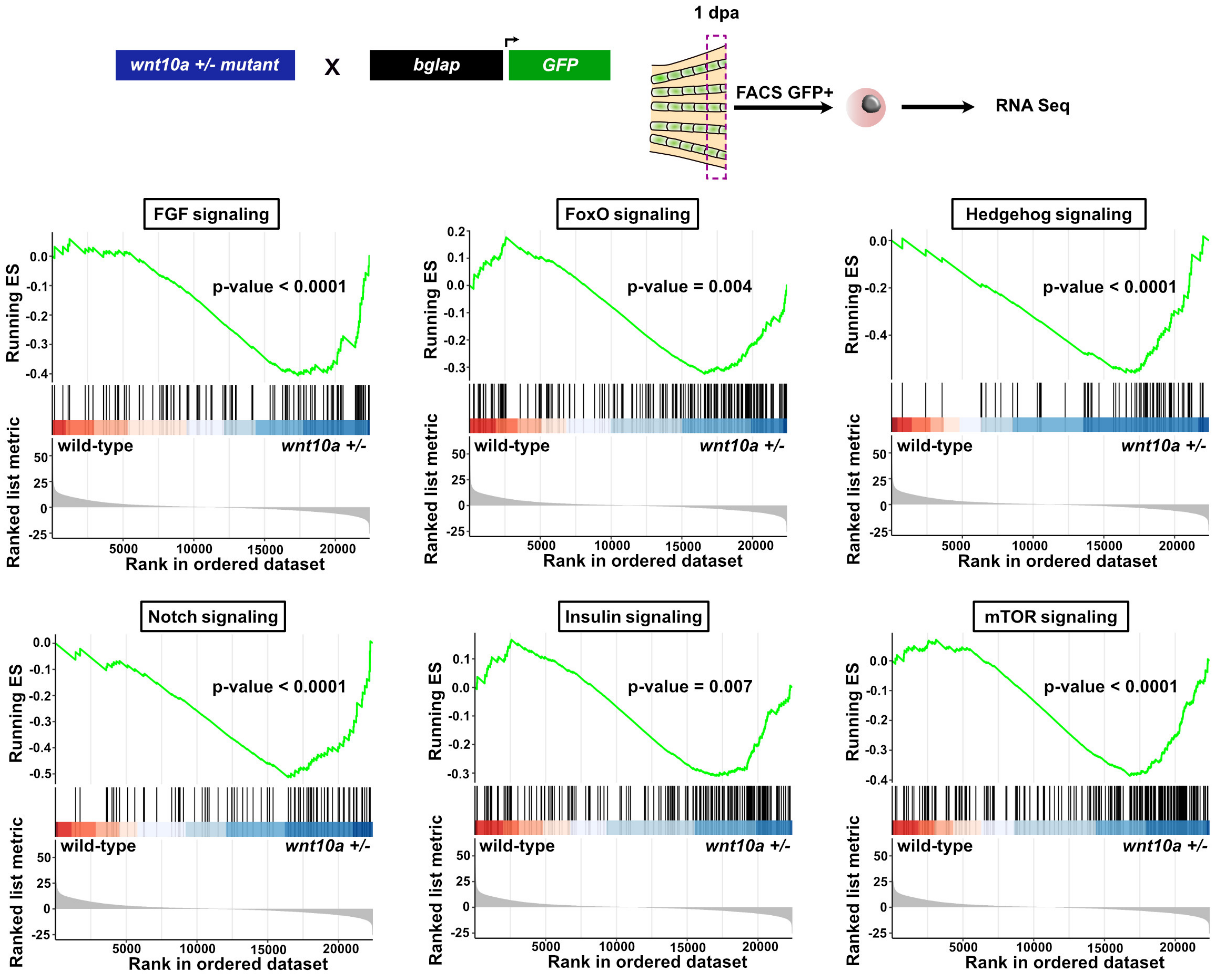


**Supplementary Figure 9.**

Running enrichment score plots show negative enrichment of genes annotated to function in several signaling pathways in RNASeq data of *bglap:GFP*+ osteoblasts isolated from *wnt10a*$^{+/-}$ fins at 1 dpa, relative to wild-type siblings.

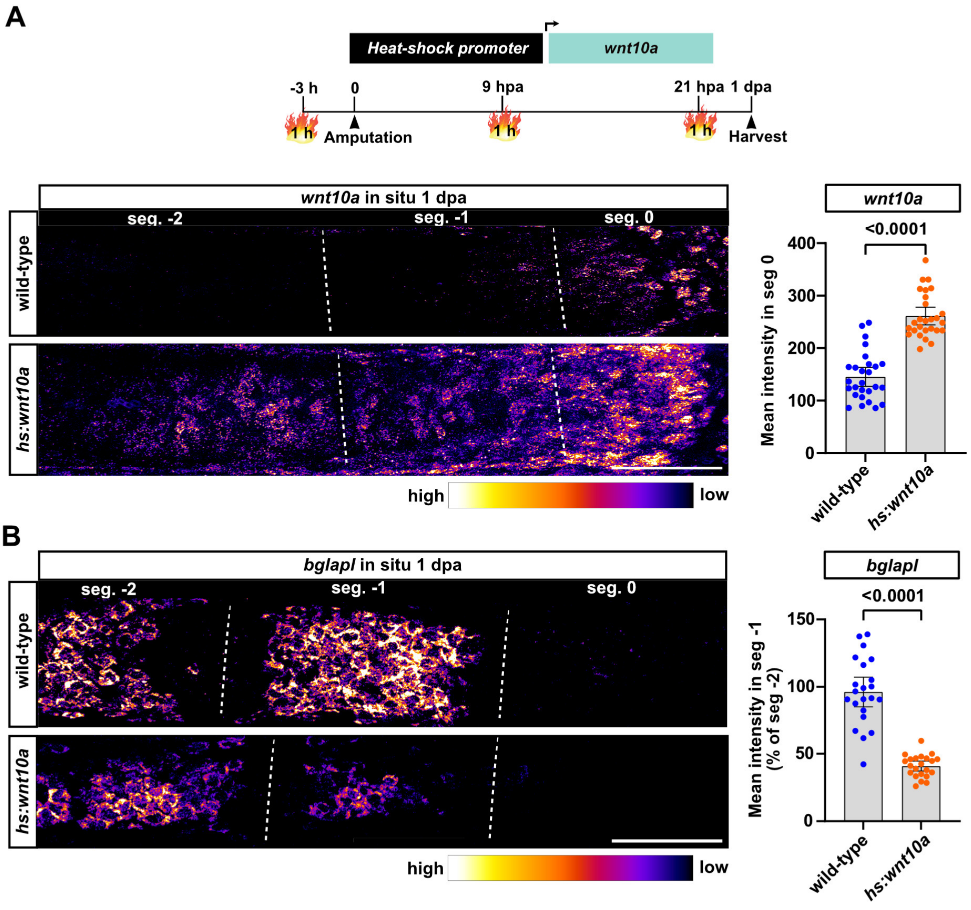


**Supplementary Figure 10.**

**A)** HCR in situ hybridization reveals higher expression of *wnt10a* in segment 0, and ectopic expression in segments -1 and -2 in *hs:wnt10a* fins subjected to the indicated heat-shock regime, relative to heat-shocked wild-type siblings at 1 dpa. $n_E = 2$, $n_A = 12$ total per group, $n_R = 24$ total per group.

**B)** HCR in situ hybridization shows increased downregulation of *bglapl* in segment -1 of *hs:wnt10a* fins at 1 dpa, indicating enhanced osteoblast dedifferentiation. $n_E = 2$, $n_A = 12$ total per group, $n_R = 22$ total per group.

**(A-B)** Data are presented as mean values. Error bars, mean ± 95% CI. Two tailed Student´s t-test. Dashed line, joints. Scale bar, 100 µm.

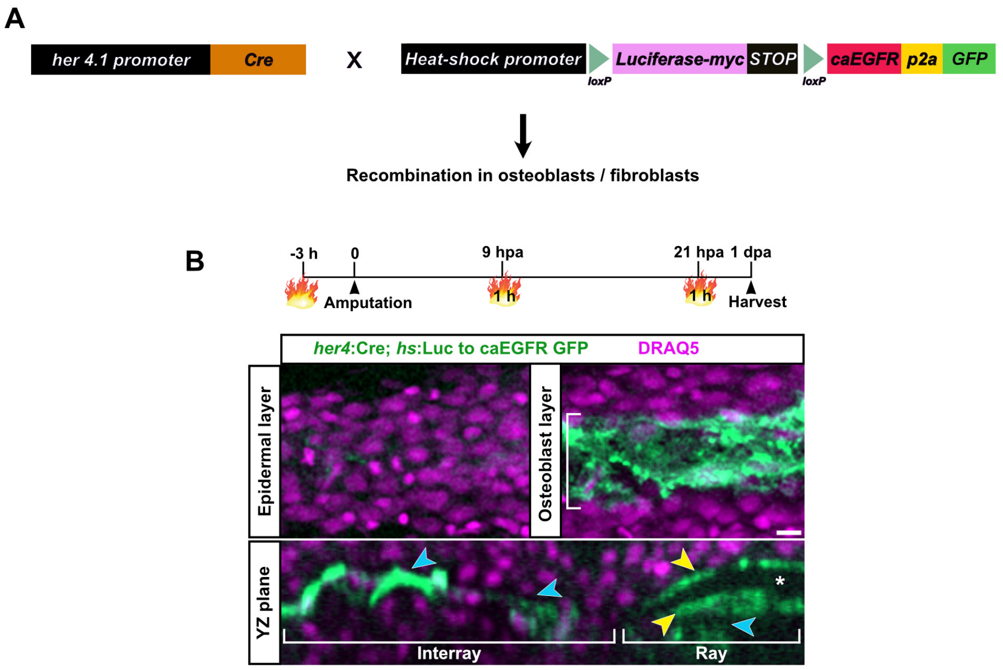


**Supplementary Figure 11.**

**(A)** Combination of the *her4.1 (3.4):Cre, cryaa:YPet*$^{ulm17Tg}$ *(*abbreviated *her4:Cre)* driver line with the heat-shock inducible Cre responder line *-1.5hsp70l:LOXP-Luciferase-MYC-STOP-LOXP-caegfra_L857R-2A-EGFP,cryaa:AmCyan*$^{ulm20Tg}$ (abbreviated *hs: Luc to caEGFR GFP*) expressing GFP (together with a constitutive-active EGF receptor) for testing the tissue-specificity of the *her4* driver line.

**(B)** Detection of native GFP fluorescence in osteoblasts lining the bony fin rays (yellow arrowheads) and in fibroblasts located in the interray and intraray mesenchyme (blue arrowheads) in double transgenic fins subjected to the indicated heat-shock regime. Asterix indicates bone. Scale bar, 10 µm.

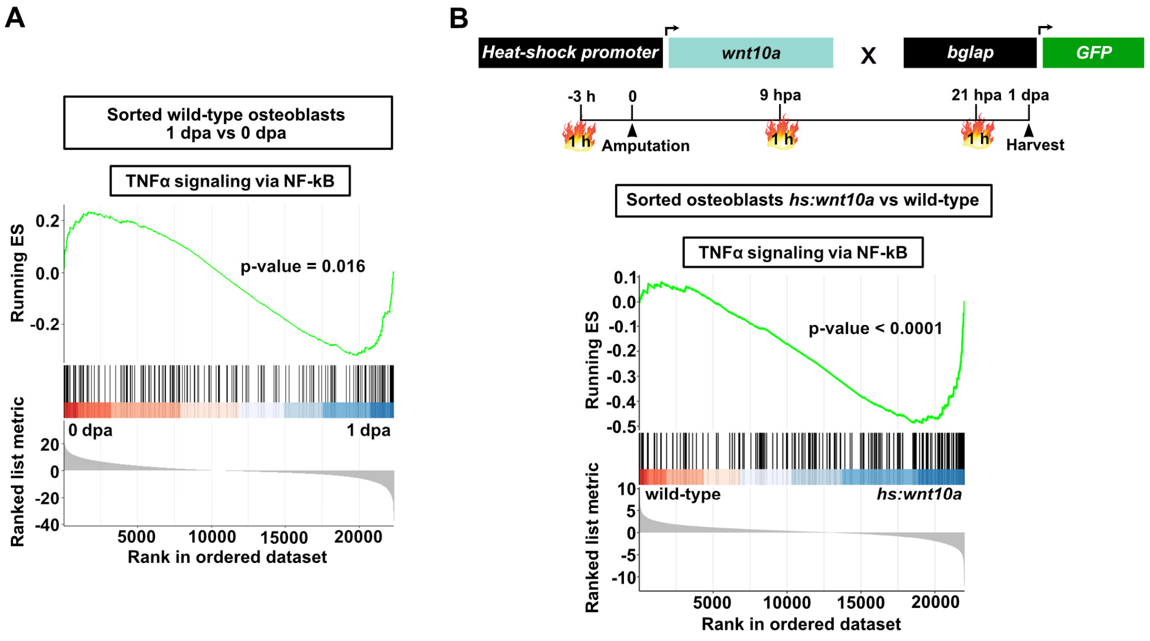


**Supplementary Figure 12.**

**A)** Running enrichment score plot shows negative enrichment of genes annotated to function in NF-κB signaling in wild-type *bglap:GFP+* dedifferentiating osteoblasts at 1 dpa relative to 0 dpa osteoblasts.

**B)** Running enrichment score plot shows negative enrichment of NF-κB signaling in osteoblasts overexpressing *wnt10a*, isolated at 1 dpa.

### Supplementary Tables

#### Supplementary Table 1. Primers for RT-qPCR on zebrafish samples

| Gene | Identifier (ZFIN) | Sense primer | Antisense primer |
|---|---|---|---|
| *actin, beta 2* | ZDB-GENE-000329-3 | ACGATGGATGGGAAGACA | AAATTGCCGCACTGGTT |
| *hatn10* | ZDB-NUCMO-180807-7 | TGAAGACAGCAGAAGTCAATG | CAGTAAACATGTCAGGCTAAATAA |
| *bglap* | ZDB-GENE-050113-4 | GCTGCAGAATCTCCTAATCATGA | TCTCCAGGTGCAGTTCCAG |
| *bglapl* | ZDB-GENE-060608-3 | AGAGTTTGCGAGAGGTGTGT | TGTAAGCCGCTACGATCCC |
| *p63* | ZDB-GENE-030819-1 | CTTATTAGGGTTGAGGGGAACAGTC | TGGTTTCCAATGTGACAATGATGAG |
| *hpdb* | ZDB-GENE-040808-62 | GCGGACAGGATATAATGGTCTCTTT | CACTATTGGCACCATCTCACTGT |
| *col1a1a* | ZDB-GENE-030131-9102 | AAACAGAACTTCCCTCGCCT | TCATTGTAGACCTGGCCGTC |
| *col10a1a* | ZDB-GENE-030131-8373 | AGTACCAGCCTTACTCCGTG | GGCAGACCTTCACCATCTTG |
| *entpd5a* | ZDB-GENE-100419-1 | AGTGGAGCTTTGGTGGACTG | TGCACAATCCCTTTCACCAC |
| *mgp* | ZDB-GENE-060928-1 | GAACCCATATCAGGCCAACG | GGCCACCTGAGAACCGTAG |
| *sp7* | ZDB-GENE-040629-2 | GACCCTCACTGGACTGCTTC | GGATGGTGCTTCCCGGTTTA |
| *sparc* | ZDB-GENE-030131-9 | TGCGGCACTAACAACAAGAC | GTCACCAGCACGTTCTTCAG |
| *bgnb* | ZDB-GENE-040426-21 | AAGTCACGCCGGTCTGAATA | AGAGCGGCAGGAGAACATC |
| *bmp8a* | ZDB-GENE-030912-13 | GTTTCAACCACACCCTCCAT | ACATCAAATGCCAGCCAACC |
| *bmper* | ZDB-GENE-030219-146 | TCCTGCCCGTCTCATCTTAC | TGGCGATAAGTCAAACACCC |
| *cadm1b* | ZDB-GENE-080327-34 | TCCGTCATCCAACTGCTCAA | GACACCAAGAGCTCGTTGTC |
| *calcrla* | ZDB-GENE-040822-26 | TAACAAGAAGGAGCGCTCGT | CAAATCGTCCAGCAGCTCG |
| *cd81b* | ZDB-GENE-040808-52 | CGTCACGACAGTCAAACCAG | CACCGATCGCTATGAGGACA |
| *ccn4b* | ZDB-GENE-080220-16 | GACCGCTTGTGGATTGTACC | AGGAGGAGAGTCTGTGGAGT |
| *clec11a* | ZDB-GENE-130530-861 | GACGCCATTCAGCAAGTCTC | TCTGCGTAGGTCTCGTACAC |
| *hhipl2* | ZDB-GENE-131120-145 | AGTGCGGTGATTTGGTAAAGG | TGTGGAAGTCTGCGCAGTA |
| *satb2* | ZDB-GENE-070912-212 | ACAAAAGCTGGGCCGTCTA | TCCTGACCAGCACAAACTCA |
| *scxa* | ZDB-GENE-060503-414 | GACCCTCCACCTCAAACTACT | GTTTAACTCCTCAGGGCGGA |
| *sema3aa* | ZDB-GENE-991209-3 | CAAAGAGAACGTGCCCAGAC | CCTCCGCTCCAACCAGTAAT |
| *sema3c* | ZDB-GENE-050513-5 | ACAGCCCAGTGTGTGCATAT | TGAGCATGACCGAGACAGTG |
| *sort1a* | ZDB-GENE-040426-2329 | ATTTACAGCAGCCGGACCAT | GGCCAAAGACACTGAACCAC |
| *tent5ab* | ZDB-GENE-071004-35 | GGCTACAAGGACCTGGATCT | ACTCCATCGGTCTGAATCGT |
| *adkb* | ZDB-GENE-030131-948 | TGATCGTAACTGGAGTCTGGT | TTGTCGGACGCGTGTTTG |
| *ankha* | ZDB-GENE-050913-33 | TTCAGTTGGAAGCAAGACCA | TTTCAGGAGGATGCCAGCA |
| *atad1b* | ZDB-GENE-030616-44 | TCCAGACTTCTTCAGCCACC | AACGGAAACCAGCCTCTTTG |
| *CABZ01111959.1* | ENSDART00000169103.2 | TGGATTCCACGAGTGTCTCA | CTTTTCCCACACTCCAGACG |
| *casp21* | ZDB-GENE-190425-1 | CCCTGTGCATTTGAGCCCTA | TGTGCTTGAAAGTCTGCGTT |
| *dph6* | ZDB-GENE-050227-15 | CATCAAAGTGGCAGCATTCG | GACAGTCGAGGGTGAAGGT |
| *hmga2* | ZDB-GENE-030131-3148 | AGGCTTCAGGATCGCAGG | AGGTCTTTTCTCCCCTGTGG |
| *il34* | ZDB-GENE-050419-150 | GGTGTTCAGACTGCGAAACA | CCTGTTGACTCACGTAAAGCC |
| *nfe2* | ZDB-GENE-030124-1 | CGGTGTACTTCCCCTGAGG | TGATGGCCATTAGTTCCTGC |
| *pea15* | ZDB-GENE-170119-2 | AGAATGACAAACTGGCGCAA | CGTGTGAGTTTGGTGTCGAT |
| *pus3* | ZDB-GENE-030131-9433 | CAATGGGCCCCTGTAGAGAG | TGAAGTCATGCGTTCCCTCA |
| *si:ch211-112c15* | ZDB-GENE-141211-78 | AAACGGAACATCTTCGCAGC | GTACGTATACTTTGGGAGGCTAT |
| *smfn* | ZDB-GENE-040426-1709 | CAAGCTCAACGGATGGTCTG | AGTCTGTAACAATGCAAGCCA |
| *syt12* | ZDB-GENE-081022-147 | AACTTCCCCGGATGCTTCAT | CAAATACCACGGTGAAGGCG |
| *them4* | ZDB-GENE-070112-982 | TCAAGTGCAATTCTACGGCG | AGTCCTGCAGCTGAGATGAA |
| *wnt10a* | ZDB-GENE-990415-278 | AACGTGGAGTATGGGGAGC | CCCGATGTACCATGGCATTT |
| *fgf20a* | ZDB-GENE-060110-1 | CGGCTACCACCTCGAGATAT | GCCATGCCGATACAGGTTAG |

| | | | |
|---|---|---|---|
| *dkk1b* | ZDB-GENE-990708-5 | AGTTTCGACTCAAGGATCACC | AGTCCCTCTGCACAATCTGA |
| *lef1* | ZDB-GENE-990714-26 | AGTATTACGAATTAGCCCGCA | GGCTTCCATCTCTAAAAGGGG |
| *axin2* | ZDB-GENE-000403-2 | CACGAGTGCCAATGACAGTG | ATCTCCTTAGGTGGCCGC |
| *NF-κB iaa* | ZDB-GENE-040426-2227 | CCCTAACTACAGCGGACAC | CAAGCTGGACCAGACTCTC |
| *NF-κB iab* | ZDB-GENE-030131-1819 | CAGACACCAAACTATAATGGTCAG | TTTATATCTGCGCCGAGCT |
| *runx2b* | ZDB-GENE-040629-4 | CTGGATGACTCTCCGAAGGC | AGCCTGTCGTGGATCTGTAA |

#### Supplementary Table 2. Hybridization chain reaction (HCR) in situ probes

| Gene | Gene Identifier | designed by |
|---|---|---|
| *bglap* | NM_001083857.3 | Molecular Instruments, Inc |
| *bglapl* | ENSDART00000171799.1 | Molecular Instruments, Inc |
| *bgnb* | NM_001001825.2 | Molecular Instruments, Inc |
| *ccn4b* | NM_001113628.1 | Molecular Instruments, Inc |
| *clec11a* | XM_691681.6 | Molecular Instruments, Inc |
| *hmga2* | NM_212680.1 | Molecular Instruments, Inc |
| *nfe2* | NM_175043.2 | Molecular Instruments, Inc |
| *sparc* | NM_001001942 | Molecular Instruments, Inc |
| *wnt10a* | NM_130980.1 | Molecular Instruments, Inc |

#### Supplementary Table 3. Primary antibodies

| Short name | Full name | Source | Identifier | Application |
|---|---|---|---|---|
| Myl7 | Rabbit polyclonal Myl7 | GeneTex | Cat# GTX128346, RRID:AB_2885759 | Immunostaining: Zebrafish cryosections |
| embMHC | Mouse monoclonal MYH7 | DSHB | Cat# N2.261, RRID:AB_531790 | Immunostaining: Zebrafish cryosections |
| Zns5 | Mouse monoclonal zns-5 | ZIRC | Cat# ZDB-ATB-081002-37, RRID:AB_10013796 | FACS live staining, Zebrafish isolated osteoblasts from fin |

#### Supplementary Table 4. Secondary antibodies

| Name | Source | Identifier | Application |
|---|---|---|---|
| Goat anti-Mouse IgG (H+L) Highly Cross-Adsorbed Secondary Antibody, Alexa Fluor 555 | Invitrogen | Cat# A21424 RRID:AB_141780 | Immunostaining |
| Goat anti-Rabbit IgG (H+L) Cross-Adsorbed Secondary Antibody, Alexa Fluor 633 | Invitrogen | Cat# A21070 RRID:AB_2535731 | Immunostaining |

#### Supplementary References

Raman, R, Antony, M, Nivelle, R, Lavergne, A, Zappia, J, Guerrero-Limón, G, Caetano da Silva, C, Kumari, P, Sojan, JM, Degueldre, C, Bahri, MA, Ostertag, A, Collet, C, Cohen-Solal, M, Plenevaux, A, Henrotin, Y, Renn, J, and Muller, M (2024). The Osteoblast Transcriptome in Developing Zebrafish Reveals Key Roles for Extracellular Matrix Proteins Col10a1a and Fbln1 in Skeletal Development and Homeostasis. **Biomolecules** 14. doi: 10.3390/biom14020139
